# ChromGene: Gene-Based Modeling of Epigenomic Data

**DOI:** 10.1101/2022.05.24.493345

**Authors:** Artur Jaroszewicz, Jason Ernst

## Abstract

**Background:** Various computational approaches have been developed to annotate epigenomes on a per-position basis by modeling combinatorial and spatial patterns within epigenomic data. However, such annotations are less suitable for gene-based analyses, in which a single annotation for each gene is desired.

**Results:** To address this, we developed ChromGene, which annotates genes based on the combinatorial and spatial patterns of multiple epigenomic marks across the gene body and flanking regions. Specifically, ChromGene models the epigenomics maps using a mixture of hidden Markov models learned *de novo*. Using ChromGene, we generated annotations for the human protein-coding genes for over 100 cell and tissue types. We characterize the different mixture components and their associated gene sets in terms of gene expression, constraint, and other gene annotations. We also characterize variation in ChromGene gene annotations across cell and tissue types.

**Conclusions:** We expect that the ChromGene method and provided annotations will be a useful resource for gene-based epigenomic analyses.

## Introduction

Genome-wide maps of epigenomic marks such as histone modifications from Chromatin Immunoprecipitation followed by high-throughput sequencing (ChIP-seq) experiments and chromatin accessibility from DNase-seq or ATAC-seq experiments provide valuable information for annotating the genome in a cell type-specific manner [1–7]. Notably, approaches have been developed for annotating the genome into ‘chromatin states’ based on the combinatorial and spatial patterns of epigenomic marks inferred *de novo* from the data. These different ‘chromatin states’ can correspond to different classes of genomic elements, including enhancers, promoters, and repressive regions [8–11]. Annotations from these methods have been used for a diverse range of applications, including understanding gene regulation and genetic variants associated with disease [2, 11, 12].

While some applications rely on per-position chromatin state annotations across the genome, other applications require gene-based annotations. Notably, transcriptomic data is predominantly used for gene-based analyses [13–15]. However, taking full advantage of epigenomic data to generate gene-based annotations is less straightforward than for per-position annotations. The challenge with gene-based annotations is that the combination of epigenomic marks will vary along a gene in a position-dependent manner. Furthermore, protein-coding genes differ vastly in gene lengths, ranging from a few hundred base pairs to over 2Mb (median 30kb) [16]. One strategy for gene-based annotations is to focus on the chromatin state at the transcription start site (TSS) [8, 17]. While such an approach is largely independent of varying gene lengths, it ignores potentially important information throughout the gene body. A simple alternative strategy would be based on averaging each mark’s signal across the entire gene [18]. However, such an approach loses information about which marks co-occur along the gene, and has the potential to be heavily confounded by gene length.

Another strategy has been to partition genes into regions and cluster the genes based on mark signal in those regions. For example, one study partitioned genes into five regions: the 500bp region upstream of the TSS, the 500bp region downstream of the TSS, and the remaining gene body into equal thirds. The study then averaged the per-mark signal within each region and applied the *k*-medoids algorithm to cluster [19]. However, such an approach is dependent on the specific choices of the partition and loses information due to averaging within each partition.

Other work has solved related, but different problems. EPIGENE used hidden Markov models (HMMs) to identify transcription units [20], but is not designed for clustering them. Hierarchical or multi-tiered HMMs have been proposed to capture broader domains, but are not designed to provide a single annotation per gene in a cell or tissue type [21–23]. (Note: Henceforth, we will use the term “cell type” instead of “cell or tissue type” for ease of presentation). EpiAlign was proposed to align chromatin states between two regions, such as genes, and identify corresponding regions [24], but is also not designed for *de novo* gene-based annotations.

There is thus a need for a principled, model-based method that can be used to generate *de novo* annotations of known genes based on the combinatorial and spatial information in data of multiple epigenomic marks. To address this, we introduce ChromGene. ChromGene uses a mixture of HMMs to model the combinatorial and spatial information of epigenomics maps throughout a gene body and flanking regions. Furthermore, ChromGene can learn a common model across multiple cell types and use it to generate per-gene annotations for each.

Here, we apply ChromGene to ChIP-seq data of histone marks and DNase-seq data from over 100 cell types and produce per-gene annotations for each one. We describe these annotations with respect to their mark emissions and relate them to gene expression data and other external data, including genes with high probability of loss-of-function (LoF) intolerance (pLI) [25]. We show that ChromGene annotations show better agreement with gene expression and stronger Gene Ontology (GO) and cancer gene set enrichments than other methods. We expect the ChromGene annotations we have produced will be a resource for gene-based epigenomic analyses, and that the methodological approach will be useful for applications to other epigenomic data.

## Results

### Overview of ChromGene method

ChromGene models the set of epigenomic data across genes with a mixture of HMMs (**Fig. 1**) by extending ChromHMM [9] (**Methods, Supp. Methods, Supp. Fig. 1**). The set of epigenomic data for each gene and its flanking region is binarized at fixed-width intervals, indicating observations of each epigenomic mark, before being input to ChromGene. The data for a given gene in a cell type is modeled as being generated by one of *M* HMMs, each with *S* hidden states. The emission distribution is modeled with a product of independent Bernoulli random variables following ChromHMM. There are no constraints on allowed transitions between states of one HMM, but transitions between states of different HMMs are not allowed. There is also an initial state distribution, or the probability of seeing a state first at the beginning of the gene flanking region, over states of the HMMs. The prior probability that a gene belongs to a mixture component corresponds to the sum of initial state probabilities of the states of the component. For given values of *M* and *S*, the parameters of the model are learned from the data by an expectation-maximization approach aiming to maximize the likelihood of the model parameters. Once a model is trained, ChromGene computes the posterior probability of each of *M* mixture component generating each gene’s data in a given cell type. For that cell type, each gene is then assigned to the mixture component for which it has the greatest posterior probability.

**Figure 1:**
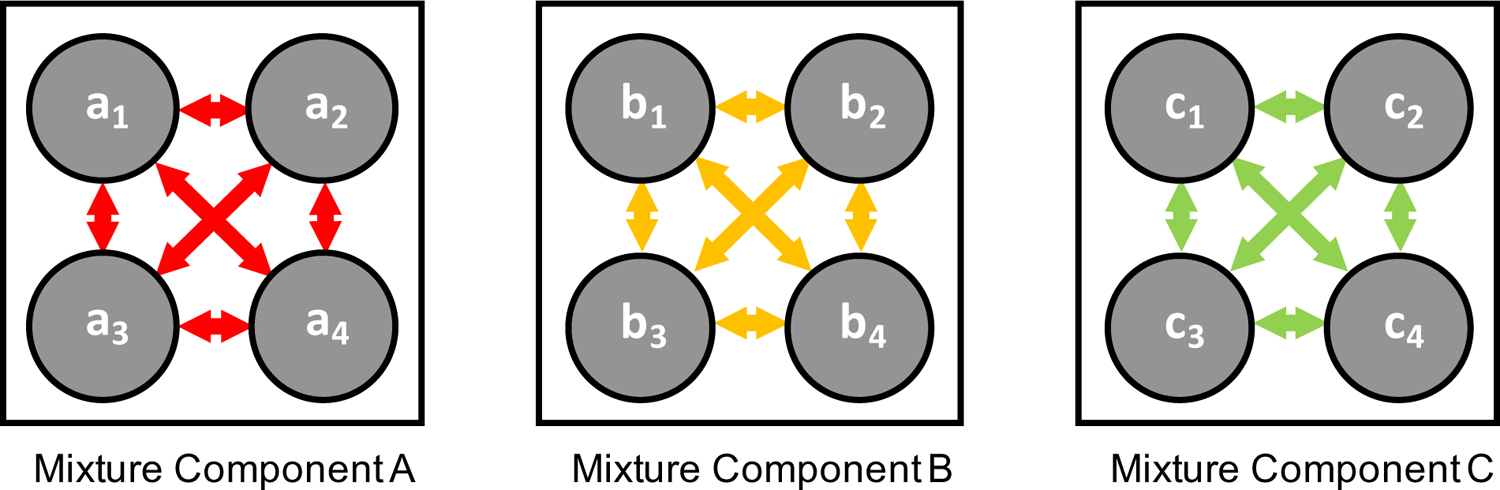
The ChromGene model Each gene is assumed to be generated by one of *M* mixture components (*A*, *B*, *C*, …), each of which is defined by *S* states. The states within a component have learned probabilities of transition between them, but transitions between states in different components are disallowed. Each state has a separate emission probability for each input mark, corresponding to the parameter of a Bernoulli random variable, and the probability of a set of observed marks is modeled using a product of those Bernoulli random variables.

To model data from multiple cell types, ChromGene can be applied analogously to the ‘concatenated’ approach of ChromHMM [2, 26], which gives cell type-specific assignments based on a common model. To do this, ChromGene treats the same gene in different cell types as if it were different genes in the same cell type when learning the model. Using this common model, a gene is independently assigned a mixture component for each cell type.

### ChromGene generates distinct gene-level chromatin annotations

We applied ChromGene to imputed data for ten histone modifications (H3K9me3, H3K36me3, H4K20me1, H3K79me2, H3K4me1, H3K27ac, H3K9ac, H3K4me3, H3K4me2, and H3K27me3), histone variant H2A.Z, and DNase-seq data from 127 cell types, binarized at 200bp resolution [1, 27] across 19,919 protein-coding genes [16] using the ‘concatenated’ approach. We focused our analysis on a model with *M=12* mixture components and *S=3* states per component to balance model interpretability while capturing biologically relevant state distinctions.

Based on the emission parameters of the model (**Fig. 2a**), along with the relationship of the components to external data not used in model learning (discussed below), we gave each component a candidate annotation (**Fig. 3, Supp. Table 1**), which we will use to refer to these components henceforth. Eight of the annotations (‘strong_trans_enh’, ‘strong_trans’, ‘trans_enh’, ‘trans_cons’, ‘trans_K36me3’, ‘trans_K79me2’, ‘weak_trans_enh’, and ‘znf’) that had at least one state with H3K36me3 or H3K79me2, both transcription associated histone modifications, present at >31% of positions. All of these annotations had at least one state associated with high frequency of promoter-associated H3K4me3 or H3K4me2 (>73%) and limited detection of the repressive mark H3K27me3 (<15%). For four of these annotations (‘strong_trans_enh’, ‘strong_trans’, ‘trans_enh’, and ‘trans_cons’), all three of its corresponding states had a high-frequency (>28%) for at least one mark. In contrast, the annotations ‘trans_K36me3’, ‘trans_K79me2’, ‘weak_trans_enh’, and ‘znf’ all had one state with low frequency of all marks (<7%). Annotation ‘znf’ is notable in that the H3K36me3 co-occurs with the repressive mark H3K9me3. This annotation shows 11-fold enrichment for Zinc finger named (ZNF) genes on average, consistent with previous findings for this combination of marks from per-chromatin state analyses [8, 27]. Three of the annotations (‘strong_trans_enh’, ‘trans_enh’, and ‘weak_trans_enh’) had a state with H3K4me1 being the mark with the greatest frequency (>78% in all cases) and limited H3K4me3 (<5% of positions), consistent with previously described putative enhancers [8, 27, 28]. Annotations ‘trans_K36me3’ and ‘trans_K79me2’ both had a state containing high frequency for active marks typically found at promoters or enhancers, a state with a low frequency of all marks, and a state dominated by H3K36me3 and H3K79me2, respectively.

**Figure 2:**
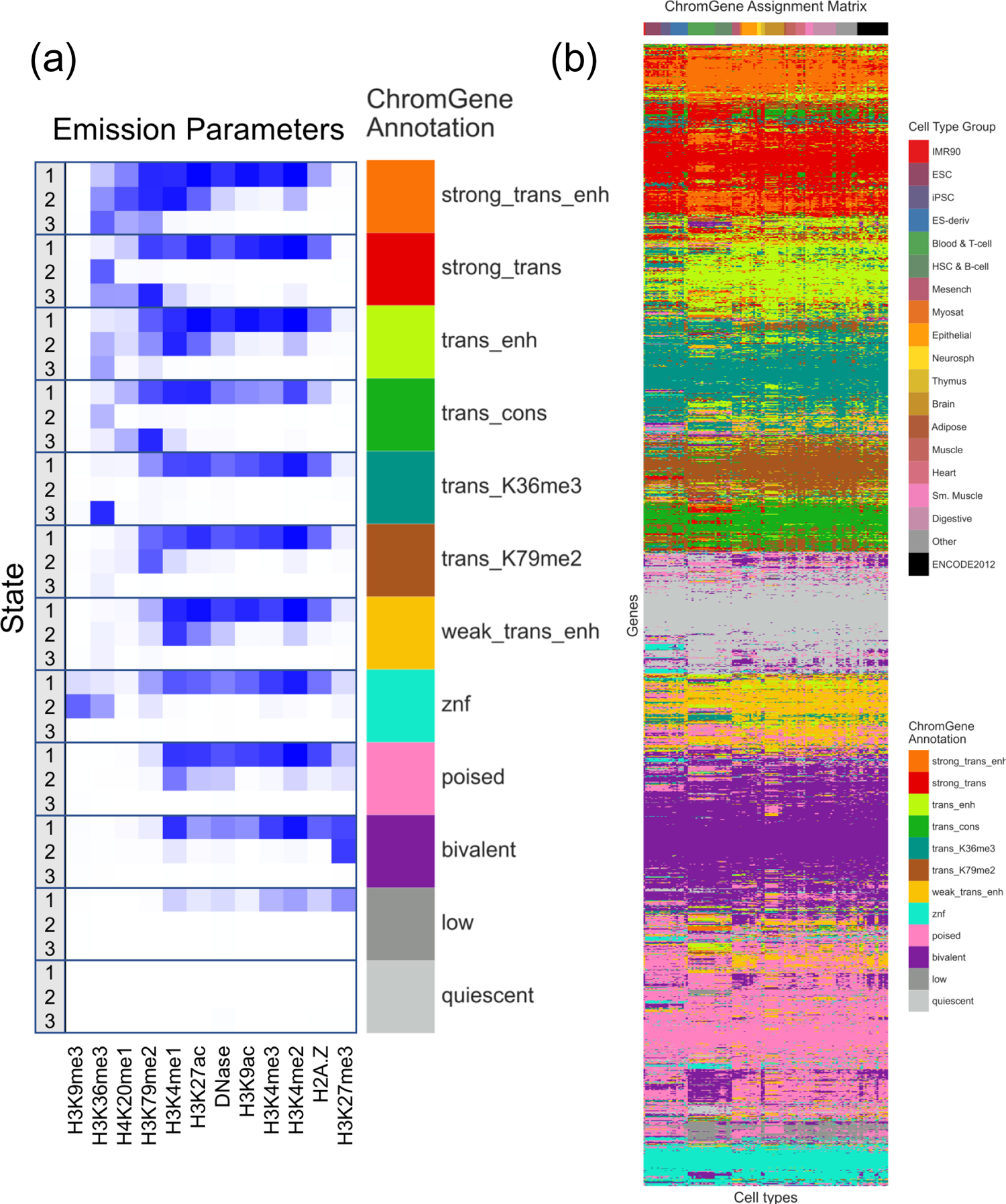
The emissions parameters and assignment of genes to ChromGene annotations. (a) Heatmap of emission parameters with blue corresponding to a higher probability and white a lower. The ChromGene annotations are labeled on the right and the states within each annotation are labeled on the left. Annotations are ordered from top to bottom by decreasing expression, and states within each annotation are ordered by decreasing enrichment at the gene TSS. Marks are ordered from left to right as previously done [27]. Transition probabilities are pictured in **Supp. Fig. 2**; per-state emission probabilities and enrichments, along with transition probabilities, are also reported (**Supp. Table 1**). (b) Graphical representation of the ChromGene assignment matrix. Columns correspond to cell types, which are ordered as previously done [1], and their tissue group is indicated by the top colorbar and top legend. Rows correspond to 2000 subsampled genes (approximately 10% of all genes). Rows were ordered by hierarchical clustering (**Supp. Methods**). Each cell is colored by ChromGene annotation for the corresponding cell type and gene (lower legend).

**Figure 3:**
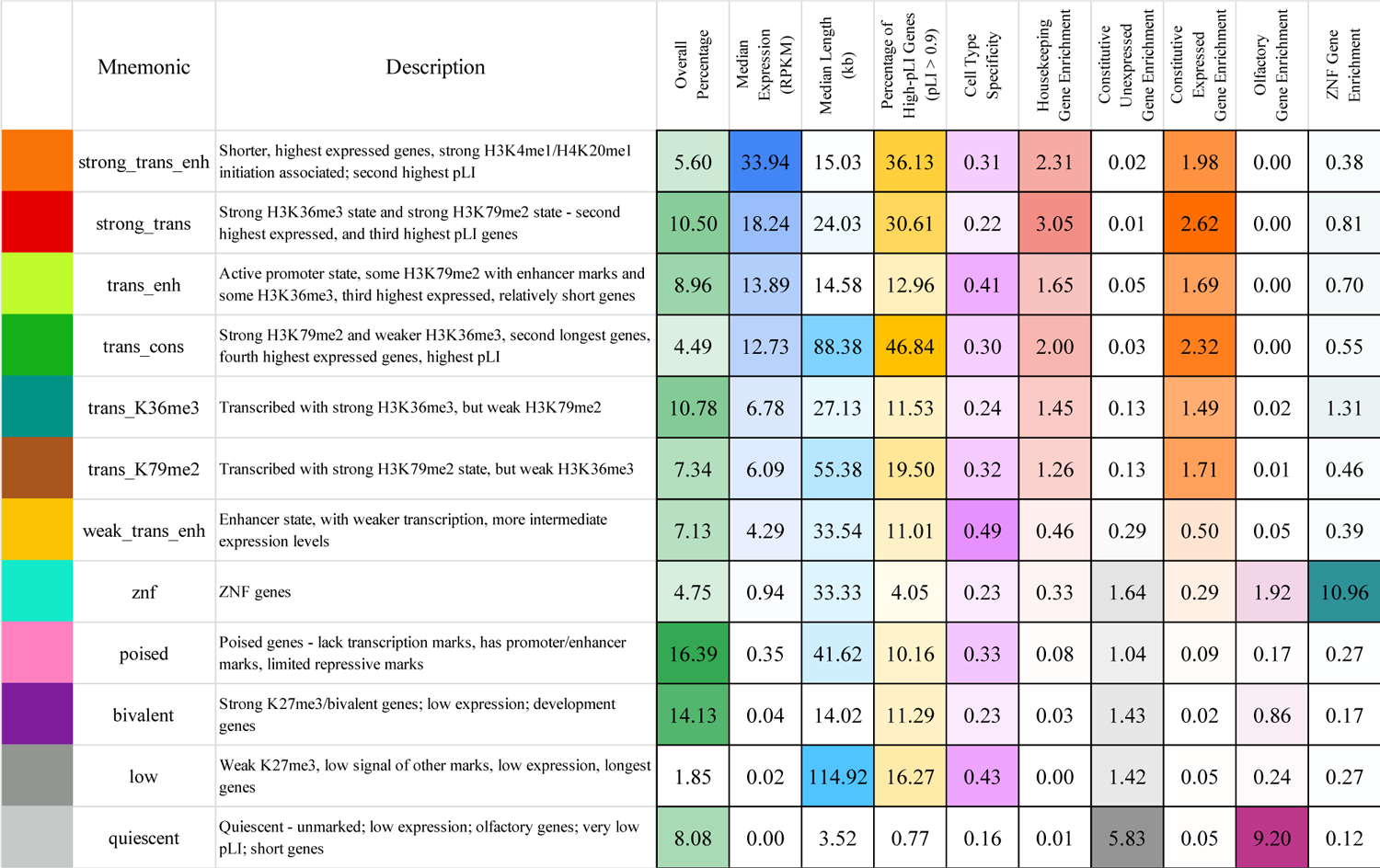
Brief description of each ChromGene annotation Rows correspond to ChromGene annotations. The columns are as follows: Color: color used for ChromGene annotation, Mnemonic: abbreviated name used for annotation, Description: description of each ChromGene annotation based on mark emissions, expression, length, pLI, and other enrichments. Subsequent columns describe summary statistics of ChromGene annotations: ‘Overall Percentage’, ‘Median Expression’ (RPKM) and ‘Median Length’ (kb) of genes assigned to annotation; ‘Percentage of High-pLI Genes (pLI > 0.9)’ – percentage of genes assigned to annotation with > 0.9 pLI (probability of Loss of function Intolerance); ‘Cell Type Specificity’ – metric of variability of annotation across cell types; ‘Housekeeping Gene Enrichment’, ‘Constitutive Unexpressed Gene Enrichment’, ‘Constitutive Expressed Gene Enrichment’, ‘Olfactory Gene Enrichment’, ‘ZNF Gene Enrichment’ – fold enrichment of gene category within annotation compared to ‘Overall Percentage’.

The four remaining annotations (‘poised’, ‘bivalent’, ‘low’, ‘quiescent’) lacked any state with the transcription associated marks, H3K36me3 or H3K79me2, at a high frequency (<12% for all states). Annotations ‘poised’ and ‘bivalent’ had states with high frequencies of other marks. Notably, the ‘bivalent’ annotation had a state associated with the high presence of the repressive mark H3K27me3 in combination with H3K4me1/2/3, along with another state associated with H3K27me3 alone. Annotation ‘low’ only had one state associated with moderate levels of histone marks, none of which were associated with transcription. No state of the ‘quiescent’ annotation had any detected modifications (all emissions <1%).

The assignments for all 127 cell types across 19,919 genes are provided as a Supplementary Data file and available at [https://github.com/ernstlab/ChromGene/]. To visualize the assignments, we sampled 2000 genes and clustered them based on assignment to ChromGene annotations across all cell types (**Fig. 2b**).

### ChromGene relationship with gene expression levels

We first investigated how the ChromGene annotation of a gene relates to its expression level. We compared ChromGene annotations to matched gene expression data for 56 cell types [1] (Methods). We separated cell type-gene combinations by their ChromGene annotation and analyzed the distribution of gene expression values (RPKM) for each annotation. We observed that genes assigned to different annotations had varying levels of expression (**Fig. 4**). The two annotations with the highest expression (‘strong_trans_enh’ and ‘strong_trans’) both had states with a high frequency of H3K36me3 and H3K79me2. The four annotations with the lowest expression (‘poised’, ‘bivalent’, ‘low’ and ‘quiescent’) each had a median RPKM of less than 1. Notably, the ‘quiescent’ and ‘bivalent’ annotations had a large overlap in distribution of gene expression levels, despite the lack of the former with any state with substantial frequency of histone modifications, and the association of the latter with histone modifications in multiple states. This highlights how ChromGene provides additional information about genes not captured by gene expression.

**Figure 4:**
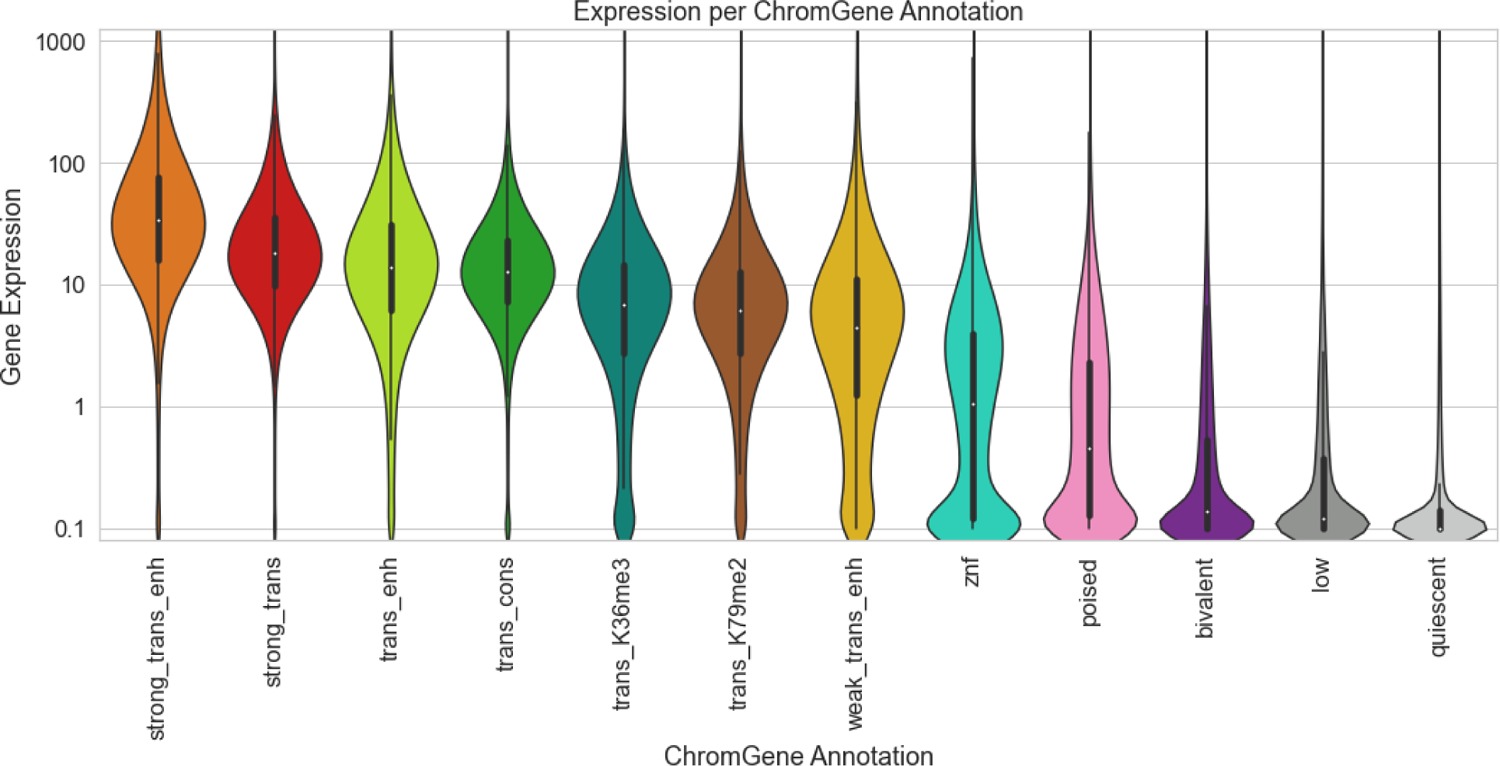
Gene expression distribution of ChromGene annotations The gene expression distribution for each ChromGene annotation across 56 cell types. Gene expression values are in RPKM after adding a pseudocount of 0.1, and then log_10_ transforming for visualization.

For each ChromGene annotation, we also determined how consistent average gene expression levels were across cell types. Specifically, for each annotation and cell type, we calculated the median log_10_(RPKM + 0.1) expression and analyzed the medians as a function of annotation (**Supp. Fig. 3**). We then calculated the mean within-annotation variance across cell types (0.026) and the total variance of the computed medians (0.783). We found that median expression variance within annotations were significantly smaller than across annotations (p-value < 10^-4^, permutation test), indicating ChromGene annotations are broadly informative of expression across cell types.

We also quantified how predictive ChromGene annotations are of gene expression and compared it to several baselines. To do this, we first separated genes into two groups: ‘expressed’ genes with RPKM ≥ 1 and ‘unexpressed’ genes with RPKM < 1. We held out genes on one chromosome at a time as a test set and used genes on all other chromosomes as a training set. We calculated the median expression for each ChromGene annotation for genes in the training set and used this as a predictor for the expression of genes in the held-out chromosome and found that the ChromGene annotation was a strong predictor of whether genes were expressed or unexpressed (across 56 cell types: mean AUROC = 0.893, standard deviation = 0.021). We also calculated the mean squared error (MSE) between the predicted log_10_(RPKM + 0.1) expression and the observed expression values across all cell types, and we found that predicted expression values were close to the true expression (across 56 cell types: mean MSE = 0.418, standard deviation = 0.050; Pearson r = 0.763). We repeated the evaluations for the following three baselines (**Supp. Methods**):

#### TSS Model

Clusters genes based only on histone mark information at the TSS. This model only takes advantage of one position, and thus does not capture additional information throughout the gene.

#### Gene Average Model

Clusters genes based on the average histone mark values throughout the gene, which is equivalent to using ChromGene with only one state per HMM. This model has two key disadvantages: First, it does not take advantage of spatial information; instead, it treats heterogeneous spatial data as if it were homogeneous, eliminating potentially useful information. Second, it is more likely to be biased by gene length because histone marks preferentially associate with specific genic regions that scale differently with gene length (see below).

#### Collapsed Model

Clusters genes using a single state per mixture component, as in the gene average model, but here the single state is found by first training a normal multi-state ChromGene model, then “collapsing” each multi-state HMM into a single-state HMM by taking the weighted average of the states in that component. This is meant to show that the differences between ChromGene and a single-state model are likely due to incorporating spatial information of histone marks instead of different instantiations of the models.

We implemented each baseline method to have an identical number of clusters as our ChromGene model. ChromGene annotations were significantly more predictive of gene expression than the three baseline methods (baseline AUROC = 0.818, 0.889, and 0.888; Pearson r = 0.601, 0.757, and 0.753; MSE = 0.685, 0.429, and 0.438; p < 10^-4^ for all comparisons, paired binomial test, for TSS, Gene Average, and Collapsed Models, respectively, **Methods**).

### ChromGene reduces association of gene length with clusters

We next analyzed the relationship between the ChromGene annotations and the lengths of genes assigned to them (**Fig. 5**). We found that the length distribution of most annotations largely overlapped, and were concentrated between 10kb and 100kb, but there were a few exceptions. Notably, ‘low’ and ‘trans_cons’ had median gene lengths around 100kb, while ‘quiescent’ was mostly composed of genes of less than 10kb length. This ‘quiescent’ annotation was strongly enriched (9.2 fold) for olfactory genes, which have a median length of 1034bp and are expressed at low levels in most cell types (mean expression=0.04 RPKM, median expression=0).

**Figure 5:**
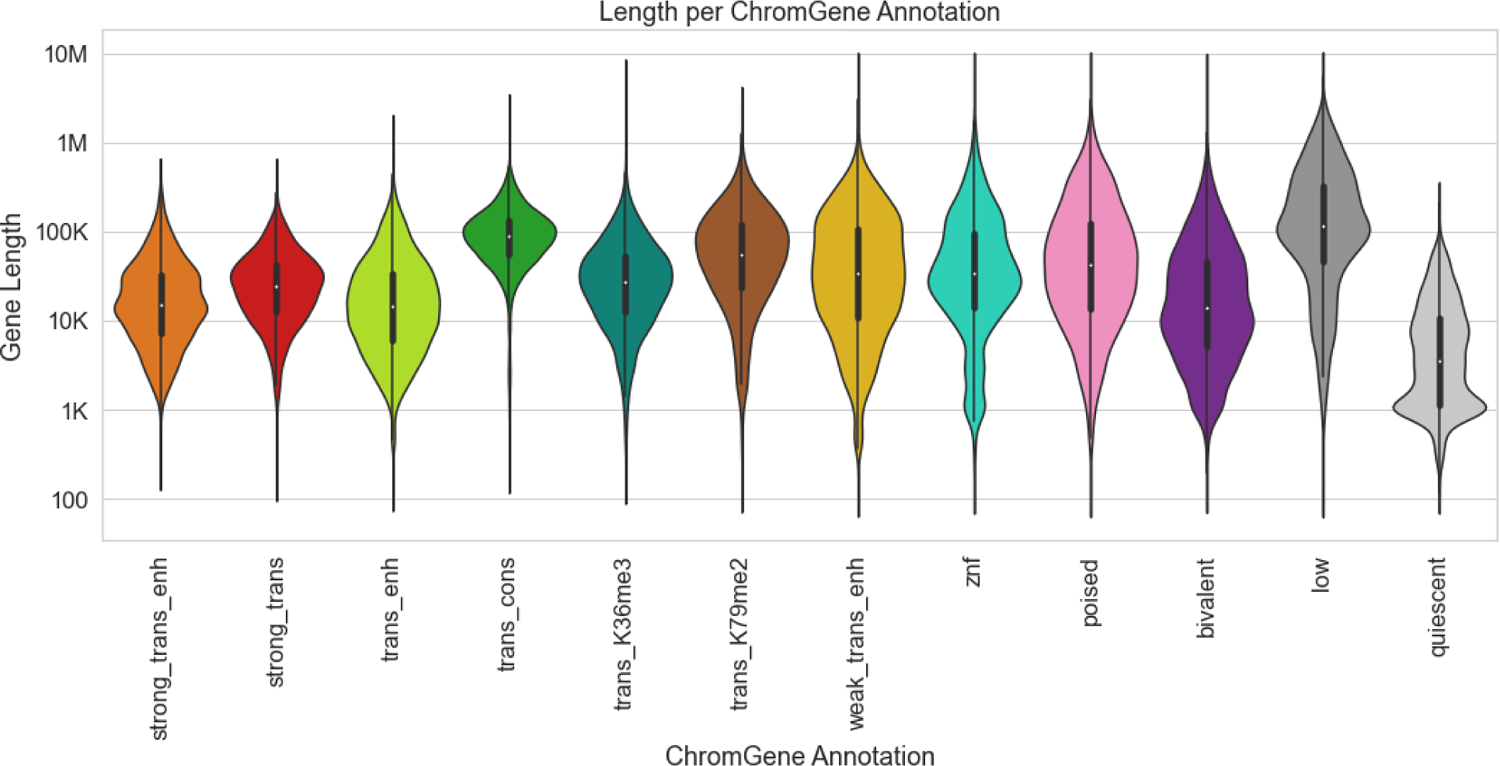
Lengths of genes by ChromGene annotation Length distribution of different ChromGene annotations. Length axis is log-transformed for visualization.

Because gene length distributions varied across ChromGene annotations, we wanted to ensure that ChromGene provided substantial information beyond gene length. We thus evaluated the amount of information shared between ChromGene annotations and gene length, and compared it to the three baselines. Specifically, for each model and each cell type, we calculated the mutual information of the gene annotation and the gene length. We expected that the baseline TSS Model would share the least information with gene length, as it only incorporates data from the TSS and not the whole gene, while among the models that incorporate data from the whole gene, ChromGene would share the least information. Indeed, we found that across cell types, the TSS model had the lowest mutual information, and ChromGene annotations had significantly lower mutual information (mean of 0.31) than the Gene Average and Collapsed Models (mean of 0.46 and 0.48, respectively, p < 10^-30^, paired binomial test, **Methods, Supp. Fig. 4**). This indicates that of models that use information throughout the gene, ChromGene annotations are least likely to simply reflect gene length.

### ChromGene annotations capture cell type-specific patterns

We next assessed how ChromGene assignments for the same gene vary across pairs of cell types. We first conducted enrichment analyses of ChromGene annotations across pairs of cell types relative to the number of genes assigned to each annotation (**Fig. 6**), excluding pairs of cell types that could be considered essentially biological replicates. As assignments of genes to different annotations across cell types could be expected based on technical variability, we also estimated a confusion matrix for pairs of “replicate” cell types (**Supp. Fig. 5a, Methods**). We found 81.1% concordance in assignments between replicate cell types, as compared to 10.2% expected by the number of genes assigned to each annotation. We next followed a similar process to calculate a contingency table for non-replicates – the probability of a given gene being assigned to one annotation for one cell type given it was assigned to another annotation in another cell type (**Supp. Fig. 5b**). We found that between pairs of randomly chosen non-replicate cell types, ChromGene annotations were 57.1% concordant, which is substantially lower than between replicates (81.1%) and substantially higher than random assignment (10.2%).

**Figure 6:**
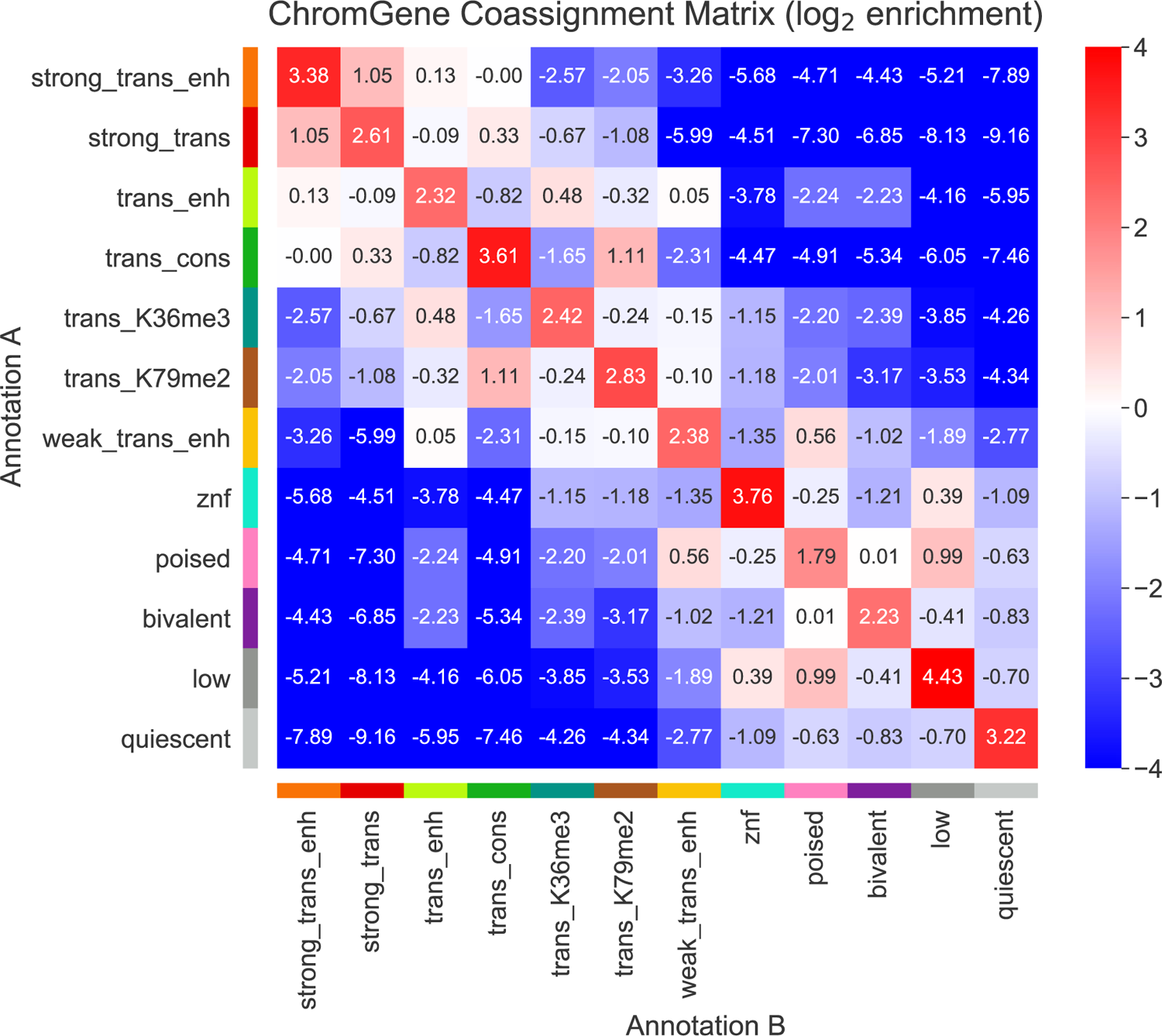
ChromGene coassignment matrix enrichment The log_2_ enrichment of combinations of ChromGene assignments over all pairs of non-replicate cell types. Enrichment corresponds to an increased likelihood of two cell types having the corresponding ChromGene assignments for a given gene, relative to randomly choosing based on ChromGene cluster sizes (**Methods**).

We further defined the “cell type specificity” of each ChromGene annotation by dividing the diagonal of the contingency table (across non-replicate cell types) by the confusion matrix diagonal (the probability biological replicates would be assigned to the same annotation), then subtracting this value from 1 to obtain a “cell type specificity score” so that higher values correspond to more cell type-specific annotations. We found that the annotations that varied the least across cell types were ‘quiescent’, ‘strong_trans’, ‘znf’, ‘bivalent’, and ‘trans_K36me3’ with cell type specificity scores from 0.16 to 0.24 (**Fig. 3, Supp. Table 1**). The most cell type-specific annotations were ‘weak_trans_enh’, ‘low’, and ‘trans_enh’ with scores of 0.49, 0.43, and 0.41 respectively. ‘Weak_trans_enh’ having the highest cell type specificity score is consistent with this annotation being primarily associated with enhancers (**Supp. Fig. 6**), which are known to be highly cell type-specific [2, 29]. The ‘low’ annotation also had high cell type specificity despite having similarly low expression as the ‘quiescent’ annotation, and was often assigned to the annotations ‘poised’ and ‘bivalent’ in other cell types.

### ChromGene annotations are differentiated by gene set enrichments

We next analyzed the enrichment of ChromGene annotations with respect to various gene sets (**Supp. Table 1**). These gene sets included zinc finger named (ZNF) genes, constitutively unexpressed genes, house-keeping genes, olfactory genes, ‘biological processes’ Gene Ontology (GO) terms, and cancer-related gene sets [1, 16, 30–34].

ZNF genes had the highest enrichment, 11.0 fold, for the ‘znf’ annotation (median p < 10^-200^, hypergeometric test, **Methods**). Constitutively unexpressed genes (RPKM < 1 in all 56 cell types with matched expression available) [1] were most enriched (5.8 fold, p < 10^-300^) in the ‘quiescent’ annotation. In contrast, a set of previously defined housekeeping genes, based on broad and constant expression levels [35], was most enriched (3.0 fold, p < 10^-300^) in the ‘strong_trans’ annotation, which was simultaneously 99% depleted for constitutively unexpressed genes. For olfactory genes [30], we observed the strongest enrichment in the ‘quiescent’ annotation (9.2 fold enrichment, p < 10^-200^), which contained 75.1% of all olfactory genes.

For the GO term enrichments [31], we calculated an adjusted enrichment p-value for each GO term gene set for each cell type and ChromGene annotation (hypergeometric test, Bonferroni corrected for the number of combinations of 127 cell types, 12 ChromGene annotations and 6036 GO terms). For each ChromGene annotation, we identified GO terms that were enriched in the majority of cell types (adjusted p < 0.01) (**Methods, Supp. Fig. 7, Supp. Table 1**). The ChromGene annotations ‘strong_trans_enh’ and ‘strong_trans’ had the greatest number of such GO terms. The most significant terms for ‘strong_trans_enh’ and ‘strong_trans’ included core regulatory and metabolic processes with terms such as ‘SRP-dependent cotranslational protein targeting to membrane’ and cytoplasmic translation’ for ‘strong_trans_enh’ and ‘mRNA splicing, via spliceosome’ and ‘mRNA processing’ for ‘strong_trans’. The ChromGene annotation ‘bivalent’ had several neuron and development related enriched GO terms, including ‘neuron differentiation’ and ‘anterior/posterior pattern specification’. In contrast, the ‘quies’ term, also associated with low expression, showed significant enrichments for terms such as ‘sensory perception of smell’, consistent with its enrichment for olfactory genes. The different GO enrichments for some annotations with similar expression levels highlights the utility of ChromGene in extracting information beyond expression.

**Figure 7:**
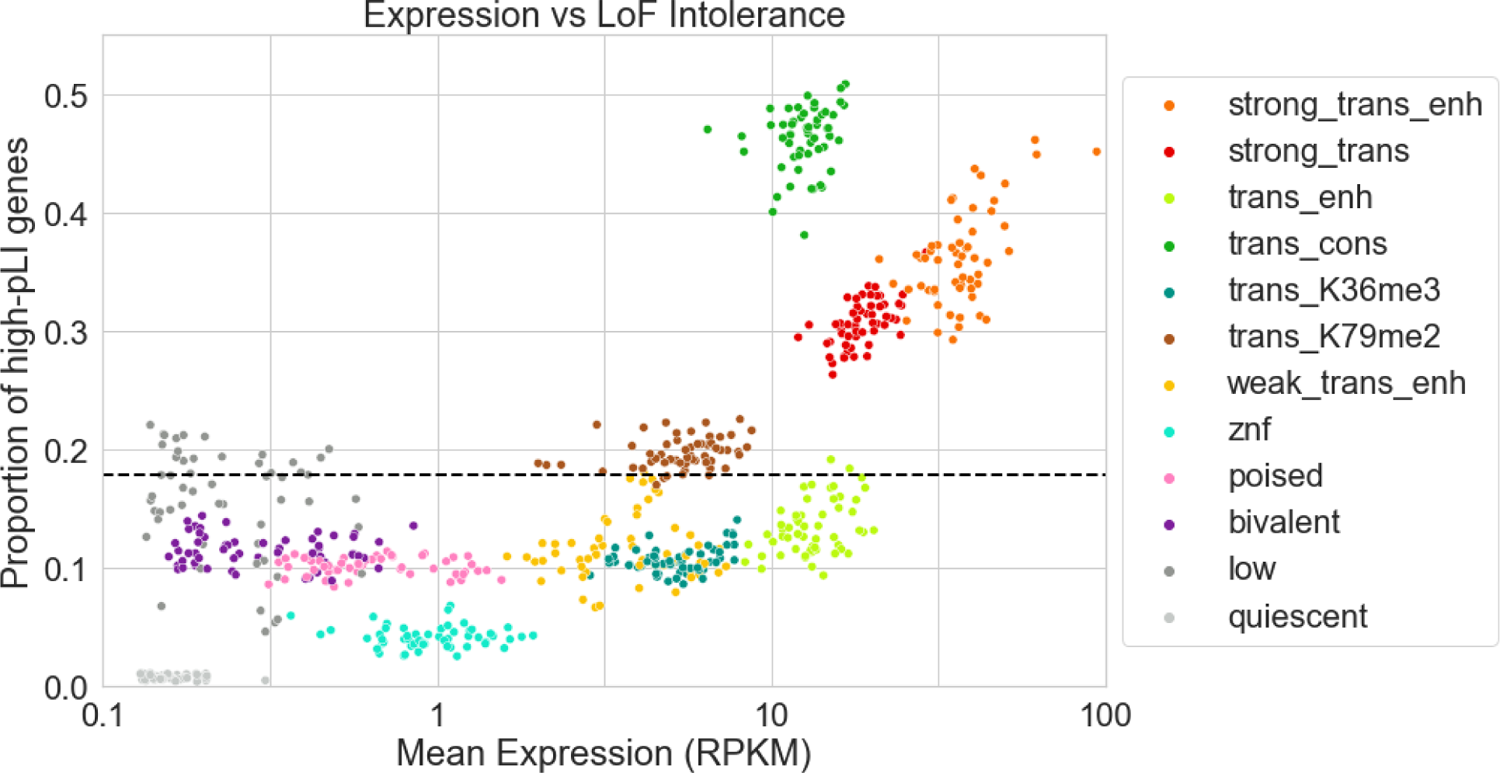
Expression and pLI scores per ChromGene annotation Scatter plot showing gene expression (RPKM) on the x-axis and the proportion of high pLI genes (pLI > 0.9) on the y-axis. Each point corresponds to genes assigned to a specific ChromGene annotation in a single cell type. Points are colored by their ChromGene annotation. Dashed line corresponds to the overall proportion of genes with high pLI. The figure shows that although there is a positive association between the two measures, ChromGene captures information beyond expression. For example, ‘trans_enh’ and ‘trans_cons’ have similar expression levels, but substantially different proportions of high-pLI genes.

Across all ‘biological process’ GO terms, we found that the ChromGene annotations ‘strong_trans_enh’ and ‘strong_trans’, which had the highest expressed genes on average, were the most likely to be enriched, constituting 26% and 44% of the significant enrichments, respectively (**Supp. Fig. 8a**). Across all ChromGene annotations, we saw an average of 133 enriched GO terms per cell type (adjusted p < 0.01). This was substantially more than for the three baseline methods (TSS, Gene Average, Collapsed), where we saw an average of 53, 109, and 82 gene sets significantly enriched (adjusted p < 0.01), respectively. These results suggest that ChromGene annotations better correspond to existing biological knowledge as reflected in ‘biological process’ GO terms than the baseline methods.

For the cancer related gene sets, we used 967 gene sets, produced by the Cancer Cell Line Encyclopedia, implicated in various types of cancer [33] and, as before, performed a gene set enrichment analysis for each ChromGene annotation. Interestingly, we found that unlike for the ‘biological process’ GO terms, the significant enrichment for cancer gene sets were most likely to be found in lowly expressed annotations, particularly ‘poised’, ‘bivalent’, and ‘low’, which constituted the majority with 21%, 60%, and 16%, respectively (**Supp. Fig. 8b**). The enrichment of the poised and bivalent annotations is consistent with previous observations that misregulation of poised and bivalent chromatin regions are associated with cancer [36, 37]. In total, we found that ChromGene had an average of 151 cancer gene sets enriched per cell type. In comparison, the baseline methods (TSS, Gene Average, Collapsed) had 71, 72, and 70 gene sets enriched (hypergeometric test, p < 0.01, Bonferroni corrected for each combination of 127 cell types, 12 annotations, and 967 gene sets), respectively, substantially less than for ChromGene. These results further support that ChromGene annotations are more consistent with established biological gene sets than the baseline approaches.

### ChromGene separates genes by pLI scores

We next explored the relationship between ChromGene annotations and genes with high pLI scores (≥ 0.9) [25]. Genes with high pLI scores have strong evidence that the gene is intolerant to loss of function, i.e., haploinsufficient.

The three ChromGene annotations most enriched for high pLI score genes were ‘trans_cons’, ‘strong_trans_enh’, and ‘strong_trans’ (**Fig. 7**). In these annotations, an average of 46.8%, 36.1%, and 30.6% of its assigned genes had high pLI scores across cell types, respectively, compared to 17.9% expected by chance. These three annotations were also among the top four most highly expressed annotations.

Notably, ‘trans_cons’ had a substantially larger percentage of high-pLI genes (46.8%) than ‘trans_enh’ (13.0%), despite a positive correlation between pLI score and expression (spearman r from 0.18 to 0.39, mean r=0.27 across cell types, all p-values < 10^-100^, **Methods**), and ‘trans_cons’ having slightly lower overall expression than ‘trans_enh’ (median RPKM = 12.73 and 13.89, respectively). This difference is consistent with the substantial difference in gene length between the ‘trans_cons’ and ‘trans_enh’ annotations (**Fig. 5**), and a previously noted positive correlation between gene length and pLI [25]. Another difference between ‘trans_cons’ and ‘trans_enh’ is that genes annotated as ‘trans_cons’ are more likely than ‘trans_enh’ to be assigned to the ‘strong_trans’ or ‘trans_K79me2’ annotations in other cell types (**Fig. 6, Supp. Fig. 5b**). Among the lowly expressed ChromGene annotations, ‘low’ and ‘quiescent’ had the greatest difference in proportion of high pLI genes (16.3% and 0.8%, respectively). Compared to the ‘quiescent’ annotation, the ‘low’ annotation contained longer genes, and genes more likely to be assigned in other cell types to ChromGene annotations with a greater proportion of high pLI genes. Similar patterns also held when considering mean pLI (**Supp. Fig. 9**). These results further show that ChromGene captures additional information beyond expression.

## Discussion

Here, we introduced ChromGene, a principled, model-based method that uses a mixture of HMMs to annotate genes based on maps of multiple epigenetic marks across genes. ChromGene’s focus on gene annotations is complementary to well established approaches for generating per-position annotations [9, 10].

We applied ChromGene to imputed data for 12 epigenomic marks to annotate genes in over 100 cell types. We showed that ChromGene annotations frequently reflect distinct gene expression levels. In cases where ChromGene annotations had similar gene expression levels, we found they differed on other important properties such as enrichment for high pLI score genes and gene set enrichments, reflecting that ChromGene annotations capture information beyond gene expression levels. We also showed that ChromGene yielded better agreement with gene expression data and more significant enrichments for cancer gene sets and GO terms than baseline approaches. We note that while ChromGene annotations will likely be preferable for many gene-centric analyses relative to per-position chromatin state annotations, it is likely less useful for when the object of study is not inherently gene centric, such as studying enrichments for GWAS identified variants.

There are multiple possible additional types of applications and extensions of ChromGene that can be investigated in future work. Although here, we applied ChromGene to protein-coding genes, ChromGene is more general and can also be applied to other pre-defined sets of genomic intervals such as long non-coding RNAs and pseudogenes. In this study, we combined our training data in a “concatenated” mode, where chromosomes across different cell types are treated as separate, and a single model is trained that applies to each epigenome. However, the input data for ChromGene can also be “stacked”, where the emissions of marks across cell types are all observed simultaneously [26, 38]. This can be used to learn patterns of variation across cell types and annotate genes based on them. In this study, we applied ChromGene on a set of 12 imputed data sets to define annotations based on a large number of cell types with a consistent set of input features. However, ChromGene can be applied with other choices for the input data.

We expect that the ChromGene annotations we have generated will be a useful resource for gene-based analysis in and across many cell types, and that the approach will be useful for gene-based annotations for data from additional marks, cell types, or species.

## Methods

### ChromGene model

For a single cell type, ChromGene uses a mixture of multivariate HMMs to model the combinatorial and spatial patterns within multiple epigenomic maps across a set of genes and to derive a single annotation per gene. Each gene is assumed to be generated by one of *M* fully-connected HMMs, each with *S* states. Each of these *M* HMMs is defined by three sets parameters: 1) initial probability for each state within each HMM, 2) the probability of transitioning from one state in the HMM to another state in the same HMM, and 3) the probability of observing each of *E* binarized emissions (in this case, histone marks or DNase mark) given a state in one of the HMMs. Although not explicitly trained, each HMM’s prior probability of being chosen for a gene is equal to the sum of the initial probabilities of its states.

More formally, we denote mixture components as *m*, *1 ≤ m ≤ M*, where *M* is the total number of components; *S* is the number of states per mixture component, and each state *s* is *1 ≤ s ≤ S*. The total number of states is thus *M*S*, and each hidden state is denoted *h_m,s_*. The initial probability, transition probabilities, and emission probabilities are defined follows:

#### Initial probabilities

The initial probability for each state, denoted τ_m,s_, corresponds to the probability that a randomly chosen gene starts in state *h_m,s_*. The sum of all the initial probabilities of states in a mixture component *m* corresponds to the initial probability of a gene starting in component *m: τ_m_ = ∑^S^ _s=1_ τ_m,s_*, and the prior probability of the gene being assigned to component *m*.

#### Transition probabilities

Each mixture component is modeled as a fully-connected HMM. The transition parameters *α_ms,ms′_* correspond to the probability of a state *s* within component *m* transitioning to a state *s’* within the same component *m*. Transitions from states within one component to states in another component are not allowed.

#### Emission probabilities

The probability of observing a specific combination of marks in a state *s* of component *m* is calculated based on the product of independent Bernoulli random variables, with emission parameters *β_m,s,e_*, over all marks following the approach of ChromHMM [9]. *β_m,s,e_* and (1 − *β_m,s,e_*) represent the probability of observing mark *e* as present or not present at a specific position in state *ℎ_m,s_*, respectively.

To apply ChromGene to multiple cell types, we apply the same modeling approach but treat data from multiple additional cell types as if they were additional genes in one cell type. This leads to a common model across cell types, but with cell type-specific gene assignments. This approach is analogous to the ‘concatenated’ model learning approach of ChromHMM [2, 26].

### Model Learning

#### Parameter initialization

The initial parameters of states across all mixture components are initialized to sum to 1 using a Dirichlet distribution with *α_m,s_* = 1. The emission parameters are initialized for each state by first randomly assigning each gene to one of the *M* components. Then, each position in the genic region and flanks is assigned uniformly at random to one of the *S* states within that component. Finally, for each of the M × S hidden states, we set the initial emission probability of each epigenomic mark to its average frequency of being observed present across all positions assigned to the state. For each component *m* and state *s* in the component, the transition parameters from the state to each state in its component are initialized to sum to 1 using a Dirichlet distribution with *α_s_* = 1.

#### Expectation-Maximization procedure

After initializing parameters, ChromGene trains the parameters iteratively with the Baum-Welch algorithm, a special case of the Expectation-Maximization (EM) algorithm, until convergence. In the E step, ChromGene takes the current parameter values (emission, transition, and initial probabilities) and calculates the log likelihood of the data being observed given those values, *log(L(θ; X, H)) = log(p(X, H|θ))*, where *θ = {θ_τ_, θ_α_, θ_β_}*, the set of initial, transition, and emission parameters, respectively, *X* is the observed data, and *H* is the set of true, but hidden state assignments. In this step, ChromGene calculates the posterior probability of each position of each gene being generated by each of the M × S hidden states.

In the M step, these posterior probabilities are used to re-estimate the parameters θ so that the new parameter values maximize the expected likelihood in the E step, i.e., *θ^t+1^ = argmax_θ_E[_H|X,θt_log(L(θ; X, H))]*. In our application, we trained for 200 iterations, as is default in ChromHMM, and parallelized the E-step computation across 8 cores. For faster computation, we also only evaluated on a randomly selected subset of the data in each iteration (**Supp. Methods**).

#### Implementation

In our implementation, we generated special input files and trained the model on top of ChromHMM [9] (**Supp. Methods**). We trained models for *M* ranging from 8 to 20, and *S* ranging from 2 to 5. To balance model interpretability while capturing major biological distinctions, we chose the model with *M=12* mixture components and *S=3* states per component for all presented results.

### Assignment of genes to mixture components

After training, ChromGene assigns each position along a gene a posterior probability of being in each of the hidden states *p(H_m,s_|X)*. From this, ChromGene calculates the posterior probability of a gene being generated by each mixture component: *P(m|X) = ∑^S^ _s=1_ p(H_m,s_|X)* based on any position. We note that all positions within a gene have the same component posterior probabilities; for example, if the first position of a gene has a posterior probability of 0.4 of being generated by component 1, then all positions within the gene have a posterior probability of 0.4 for component 1. This is a consequence of transitions between components being disallowed. ChromGene assigns each gene to the component 1 ≤ *m*^∗^ ≤ *M* with the highest total posterior probability of generating the gene, i.e., *m^∗^ = argmax_m∈M_ P(m|X)*. We used these hard assignments of genes to components for all presented results.

### Training data

To generate input data for ChromGene, we first defined our genes of interest. For this application, we used 19,919 protein-coding genes as defined by Ensembl v65 / GENCODE v10 for hg19, as the epigenomic data we used had matching gene-level expression estimates across 56 cell types based on these gene annotations [1, 16]. For each gene, we took the 5’-most base of the first exon and 3’-most base of the last exon as the TSS and TES of the gene, respectively. Genes on the negative strand were reversed to align with genes on the positive strand so all genes had the same orientation in the model. We rounded the TSS upstream (in the 3’ direction) and the TES downstream (in the 5’ direction) to the next position divisible by 200bp. We added an additional overhang of 2kb upstream of the first exon and downstream of the last exon to capture additional spatial information around the TSS and TES. We then binned the entire region (gene and overhangs) into 200bp bins so that the boundaries of each bin were divisible by 200. We then extracted the epigenomic marks, as described next.

To generate a single ChromGene model that would be comparable across a large number of cell types and marks, we used imputed data for a set of 12 marks (H3K36me3, H3K4me1, H3K4me2, H3K4me3, H3K9me3, H3K27ac, H3K27me3, H3K9ac, H3K79me2, H4K20me1, H2A.Z, and DNase) across 127 reference epigenomes, which for our purposes we treated and referred to as different cell types, except for calculation of the confusion matrix and contingency table, as described below [1, 27]. This imputed data was previously used to train a 25-state ChromHMM model. The use of imputed data allowed us to annotate more cell types with the same set of marks compared to using directly observed data. We used the same binarization for the imputed data as previously generated [27] [Data available at https://egg2.wustl.edu/roadmap/data/byFileType/chromhmmSegmentations/binaryChmmInput/imputed12marks/binaryData/].

### Gene expression data

We downloaded gene expression data for 56 cell types from the Roadmap Epigenomics Consortium [1] [available at https://egg2.wustl.edu/roadmap/data/byDataType/rna/expression/57epigenomes.RPKM.pc.gz]. We took the provided RPKM values, added a pseudocount of 0.1, and took the log_10_ transform of the value.

### Gene expression analysis

To determine whether median gene expression variance within annotations was significantly smaller than across annotations, we performed a permutation test. Each median log_10_(RPKM+0.1) expression value for a cell type was randomly assigned to one of the ChromGene components, while maintaining 56 cell types per ChromGene component. We then calculated the mean within-annotation variance of the median expression values for each permutation and for the true ChromGene annotation. To compute a p-value, we counted the fraction of 10,000 permutations in which the mean variance value based on the permuted values was less than that of the true ChromGene annotations.

To assess if there was a significant difference in ChromGene’s AUROCs at predicting expressed genes to those of the baseline methods, we used a one-sided “paired binomial test”, where we counted how often the AUROC for each cell type for ChromGene was higher than that of the baseline method. Based on these counts we calculated a p-value, where the null hypothesis is that the two methods are equally likely to produce the higher of the AUROCs (p=0.5), and n=56 cell types with expression data as observations.

### Calculation of mutual information of assignment given gene length

To calculate mutual information of gene assignments given gene lengths, for each gene, we first calculated its log_10_(*length*), which ranged from 2.06 to 6.73, where *length* is in bp. We binned the log_10_(*length*) into 0.05-increment intervals to create a discretized distribution. Then, for each cell type, we calculated the mutual information of ChromGene assignment and the length bins. We repeated the procedure for each baseline method. To compare to the baseline methods, we used the paired binomial test described above, using mutual information instead of AUROC and n=127 cell types.

### Calculation of confusion matrix and contingency table

To generate a confusion matrix (**Supp. Fig. 5a**), we took our matrix of ChromGene assignments with entries *m_g,c_*, where each of *g=1,…,19919* corresponds to a gene and each of *c=1,…,127* corresponds to a cell type (**Fig. 2**, genes subsampled and ordered for visualization). For each gene *g*, we calculated the conditional probability *P(m_g,c_ | m_g,c’_)*, where *c* ≠ *c′* correspond to pairs of epigenomes that were originally annotated as the same cell type but of different individuals (“Rectal Mucosa Donor 29” and “Rectal Mucosa Donor 31”, “Foreskin Fibroblast Primary Cells skin01” and “Foreskin Fibroblast Primary Cells skin02”, “Foreskin Melanocyte Primary Cells skin01” and “Foreskin Melanocyte Primary Cells skin03”, “Foreskin Keratinocyte Primary Cells skin02” and “Foreskin Keratinocyte Primary Cells skin03”, “Skeletal Muscle Female” and “Skeletal Muscle Male”, “Primary hematopoietic stem cells G-CSF-mobilized Male” and “Primary hematopoietic stem cells G-CSF-mobilized Female”, “Fetal Brain Female” and “Fetal Brain Male”). In short, we sought to answer the question: “*given an entry in the assignment matrix is **mixture component i**, what is the probability that the same gene is assigned to **component j** in a replicate cell type?”* We then calculated overall conditional probabilities by averaging over all genes *g*. We represented these conditional probabilities so that the component conditioned on corresponds to a row, which implies that each row sums to 1.

To generate the gene contingency table (**Supp. Fig. 5b**), we performed the same process as for the confusion matrix, but instead of calculating probabilities for pairs of “replicate” cell types, we calculated them for non-replicate cell types.

### Cell type specificity

We defined the “cell type specificity” of each ChromGene annotation by dividing the diagonal of the contingency table (across non-replicate cell types) by the confusion matrix diagonal, then subtracting this value from 1 to obtain a “cell type specificity score” so that higher values correspond to more cell type-specific annotations.

### Calculation of coassignment matrix enrichment

To generate the coassignment matrix enrichment (**Fig. 6**), we first generated an expected coassignment matrix, where we assumed independence between assignments. For each ChromGene annotation, *m*, we calculated an empirical prior probability of observing the annotation assignment for a random cell type and gene *π_m_ = P(m) = ∑^127^_c=1_ ∑^19919^_g=1_ I(m_g,c_ = m) / (127 × 19919)*, where *I(m_g,c_ = m)* denotes the indicator function applied to the ChromGene assignment for gene *g* in cell type *c* being *m*. We took these prior probabilities and calculated an expected coassignment matrix, where each entry *(i, j)* was found by multiplying the prior probabilities of mixture components *m_i_* and *m_j_*: *P(m_i_, m_j_) = P(m_i_)×P(m_j_)*. We calculated the observed coassignment matrix by counting the frequency with which a gene was assigned to component *m_i_* in one cell type and *m_j_* in the other, averaging over all genes and all pairs of cell types that were not considered replicates. We then normalized this matrix by the sum of its values to form a probability distribution. To calculate enrichments, we divided the observed coassignment matrix by the expected coassignment matrix. Finally, we took the log_2_ of these values to show enrichments and depletions.

### Olfactory and housekeeping gene annotations

We downloaded olfactory [30] and housekeeping gene annotations [35]. We matched the gene names to the GENCODE annotation [16] to label each gene as olfactory or not olfactory, and housekeeping or not housekeeping.

### Gene set enrichments

To calculate the fold enrichments for ZNF named genes, housekeeping genes, constitutively unexpressed genes, and olfactory genes, for an annotation, we divided the mean proportion of genes from the set assigned to the annotation across all cell types by the empirical prior probability of a gene being assigned to the annotation. To calculate a median p-value for a gene set and annotation across cell types, we took each cell type and calculated a p-value using a hypergeometric test (scipy.stats.hypergeom, with X=[number of genes in annotation and gene set], M=[total number of genes], n=[total number of genes in gene set], N=[number of genes in annotation]), and then took the median of those values.

To calculate significant gene set enrichments for ‘biological process’ GO terms [31], we first found the overlap of a gene set with each of the ChromGene annotations (or baseline method) for a given cell type. Next, we calculated p-values using a hypergeometric test, as for the individual gene sets above, and corrected for multiple testing with a Bonferroni correction that controls for the number of combinations of cell types, mixture components, and GO terms tested. We repeated the process for each baseline method. We then followed the same procedure for cancer gene sets [33]. The gene sets for ‘biological process’ GO terms and cancer gene sets were downloaded from the Enrichr database [34].

To generate the GO term enrichment median p-value heatmap (**Supp. Fig. 7**), we followed the same procedure. We removed rows where the median adjusted p-value was greater than our significance threshold of 0.01, and clustered rows using ‘seaborn.clustermap’, which uses a Euclidean metric and ‘average’ linking. Finally, we -log_10_ transformed the p-values for visualization.

### ChromHMM state enrichment

For each ChromGene mixture component, we took all genes and cell types assigned to the component, and counted the total observed counts of each ChromHMM state in a previously described 25-state model [27], thus yielding a matrix of *M* ChromGene components by 25 ChromHMM states. We then added a pseudocount of 1 to these counts and normalized the rows (ChromGene components) to unit probability. We divided these probabilities by the genome-wide ChromHMM state assignment proportions across all cell types to generate an enrichment, and finally, calculated the log_2_ of these enrichments (**Supp. Fig. 6**).

### pLI score analysis

For pLI score analysis, we obtained pLI scores for each gene from gnomAD [25] [https://storage.googleapis.com/gnomad-public/release/2.1.1/constraint/gnomad.v2.1.1.lof_metrics.by_gene.txt.bgz, “exac_pLI” column]. For correlating gene pLI scores with gene expression, for each of the 56 cell types with expression data available [1], we correlated the expression value of genes with their corresponding pLI scores using spearman correlation.

## Declarations

### Ethics approval and consent to participate

Not applicable

### Consent for Publication

Not applicable

### Availability of Data and Materials

The ChromGene annotations generated in this study and the ChromGene software are available in the ErnstLab ChromGene GitHub repository, https://github.com/ernstlab/ChromGene. ChromGene annotations are also available as a Supplementary Data file.

## Authors’ Contributions

AJ and JE developed the method. AJ implemented the method and performed all analyses presented. AJ and JE wrote the manuscript. Both authors read and approved the final manuscript.

## Acknowledgements

We thank Huiling Huang for conducting some preliminary analyses of the results of the method. We also thank Adriana Arneson and Petko Fiziev, and other past and current members of the Ernst Lab, for their feedback on this work.

## Supplementary Methods

### Implementation on top of ChromHMM

To implement ChromGene on top of ChromHMM, we first observed that a mixture of HMMs can be defined in terms of a single HMM with certain transitions disallowed. Specifically, given a set of *M* fully-connected HMMs, each acting as a mixture component, this can be represented with a single, larger HMM with the addition of a “dummy state” and “dummy mark.” In the single HMM, each non-dummy state of the HMM can transition into and out of other states within its component and a single “dummy state”, but not directly to states of other HMM components (**Supp. Fig. 1a,b**). From the “dummy state”, transitioning to any component is allowed, but only through the use of the “dummy mark”, which forces transitions into and out of the “dummy state”.

We structured the input data so that all genes in a chromosome and cell type (after adding 2kb overhangs and reversing genes on the negative strand) were concatenated head to tail, but with a single “dummy position” separating the genes, and a dummy position at the beginning and end of the data. In this dummy position, the emission for all the original input marks is set to 0, but we include an emission of 1 for a single, new mark designated the “dummy mark” (**Supp. Fig. 1c**). This dummy mark has an emission of 0 within all extended genic regions, and 1 only for the dummy position (**Supp. Fig. 1d**). Upon reaching the dummy position, the model is forced to transition to the dummy state. After this dummy state, the hidden state can transition to any of the mixture components.

This structure also allowed us to compactly represent all genes without having a separate file for each gene. For a single cell type, we had 23 chromosome files (chromosomes 1-22 and X). For 127 cell types, this yielded 127 × 23 = 2921 total input data files.

Similarly, we generated initial parameter files for the initial parameters described in the Methods section. For *M* mixture components, *S* states per component, and the dummy state, we had a total of (M × S) + 1 states. The initial probability for each state in each component is set to 0, except for the dummy state, which is set to 1. The transition probabilities from any state in a component to each other state in the same component was allowed to be nonzero, and normalized so that the sum of these probabilities was 0.95. The remaining 0.05 probability was set to be the probability of transitioning from the component’s states to the dummy state. The probability of transitions from the dummy state to each of the other M × S states corresponds to the initial parameters, which, as described (**Methods**), were initialized using a Dirichlet distribution with α_m,s_ = 1 for all components so that they summed to 1. The transition probability from a state in a component to each state in different components was set to 0.

Finally, the emission probabilities were initialized as described in the Methods section, except that for the dummy state, the emission probability of the epigenomic marks was set to 0, and the emission of the dummy mark was set to 1; for remaining genomic states, the emission probabilities for the dummy mark was set to 0, and probabilities for the epigenomic marks were not restricted.

We passed the data and the initialized parameter files into ChromHMM’s LearnModel function and used several optional flags. We used the -e 0 and -t 0 flags to enforce that a 0 or 1 emission or transition parameter in the initial parameter file remains at that value. We also used the -scalebeta flag to increase numerical stability. With the -scalebeta flag, ChromHMM rescales the backward variables from the forward-backward procedure based on the sum of their values at a position as opposed to reusing the scaling value for the forward variables. We observed that this was necessary in some cases to prevent overflow in this setting. To then prevent underflow with the -scalebeta flag, if a backward variable value fell below 10^-300, ChromHMM set it to 10^-300. Similarly for a forward variable, if its value fell below 10^-300, ChromHMM set it to 10^-300, except if the emission probability product for the state at the position was 0, in which case ChromHMM kept the forward variable as is. We used the flag -n 100 to sample 100 files for each iteration, and -d - 1 to allow the log likelihood to increase between iterations. The log likelihood can increase between iterations because different subsamples of the data are evaluated on different iterations. We trained for 200 iterations, as default in ChromHMM, using ChromHMM v1.18.

After training the model, we used ChromHMM to generate posterior probability assignment files from which we calculated the posterior probabilities of each gene in each cell type being generated by each of the *M* HMMs and computed the maximum to derive hard assignments.

### ChromGene annotation visualization

To visualize ChromGene annotations across cell types and genes, we first randomly subsampled the 19,919 protein-coding genes to 2000 genes. For the ChromGene annotations for these 2000 genes, we calculated a pairwise distance between them as *(1 - mean(I(X_i_ = Y_i_))*, where *X_i_* and *Y_i_* corresponded to the ChromGene assignments across the 127 cell types and *I(x)* is the indicator function. Columns, which correspond to cell types, were ordered the same way as previously [1]. Rows were ordered using scipy’s hierarchical clustering function, using optimal leaf ordering and average (UPGMA) linkage.

### Baseline model implementations

#### TSS model

To generate a 12-state model based only on gene TSSs, we took histone mark binarization data across the 127 cell types for the 19,919 genes, and used the data at the TSS position, rounded down to the closest 200bp bin. As with the standard ChromGene input data, we separated these individual TSS positions by a single dummy position, then trained a 12-state model on top of ChromHMM and followed the same procedures for generating annotations.

#### Gene average model

The gene average model was implemented using ChromGene, but it used a single hidden state per mixture component (i.e., *M*=12 components, *S*=1 states per component). All other procedures were identical.

#### Collapsed model

The collapsed model was identical in structure to the gene average model (*M*=12 mixture components, *S*=1 states per component), but was not trained directly. After training the full ChromGene model (*M*=12 components, *S*=3 states per component), as is used in all results presented, we calculated the prior probability of a collapsed component by summing the initial probabilities within its corresponding component of the full model. If the initial probabilities for the *S* states within a component *m* are denoted τ_m,s_, then the initial probability for the collapsed state *s’* of the corresponding component *m’* is equal to the prior probability of component *m*: τ_s′_ = π_m_ = ∑^S^ _1_ τ_m,s_. The transition probability from the dummy state to a component is this prior probability. The transition probability of the collapsed state to itself is calculated by comparing the number of observed state assignments for the corresponding component in the full model to the number of genes assigned to that component. If the number of 200bp positions assigned to some component is denoted *x_m_*, and the number of genes assigned to that component is denoted *n_m_*, then there are *(x_m_-n_m_)* transitions from the component to itself and *n_m_* transitions from the component to the dummy state, for a total of *x_m_* transitions. We set the transition probability of the collapsed state to itself as *(x_m_ − n_m_)/x_m_* and the transition probability of the collapsed state to the dummy state as *n_m_/x_m_*.

The emission probability parameters of the collapsed model are set to the weighted mean of the probabilities of the full model, weighted by the state prior probabilities. If the emission probability for state *s* in component *m* for mark *e* is denoted β_m,s,e_, the emission probability of the corresponding collapsed component *m’* with the single collapsed state *s’* is β_m′,s′,e_ = _1_ π_m,s_β_m,s,e_, where π_m,s_ is the prior probability of state *s* in component *m*, which is equal to the fraction of 200bp bins assigned to state *s* across all cell types and genes. These parameters are all set in the ChromHMM model file and are used directly without training to generate assignments for all genes and cell types.

## Supplementary Figures

**Supplementary Figure 1:**
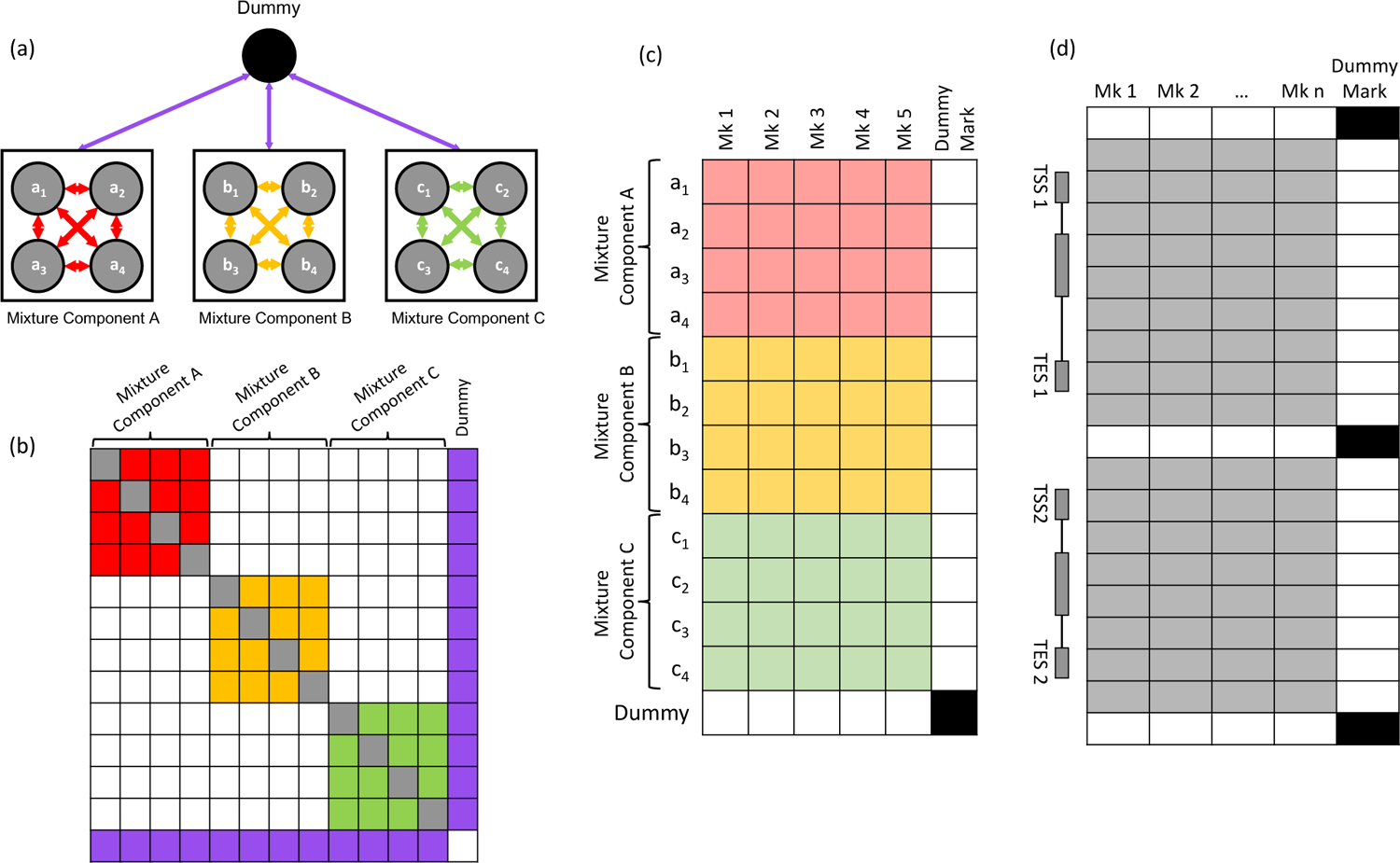
Schematic of ChromGene implementation on top of ChromHMM (a) Diagram showing *M*=3 ChromGene mixture components (*S*=*4* states per component in each black box), connected by the “dummy state”, which is used for implementation on top of ChromHMM. (b) Transition matrix for example model. Transitions are allowed within each component (within red, within yellow, within green), but transitions are not allowed directly between components. Transitions are allowed from any component to the dummy state, or from the dummy state to any components, but these transitions only occur at the single “dummy position” between genes. (c) Example emission matrix. All states within components A, B, and C (denoted a_1_…c_4_) have an emission probability between 0 and 1 (light red, yellow, and green) for the five marks, and an emission probability enforced to be 0 (white) for the “dummy mark” (last column). The “dummy state” (last row) has an emission probability enforced to be 0 for all marks except for the “dummy mark”, for which the emission probability is set to 1 (black). (d) Example input data. At the dummy position, only the dummy mark is emitted (right column, black), and no other marks are emitted at that position (white). Within the gene body, epigenomic marks may be emitted (grey), but the dummy mark cannot (white). Only the dummy state can emit the dummy mark. This enforces that the dummy state is observed between genes to allow sequential genes to be assigned to different components.

**Supplementary Figure 2:**
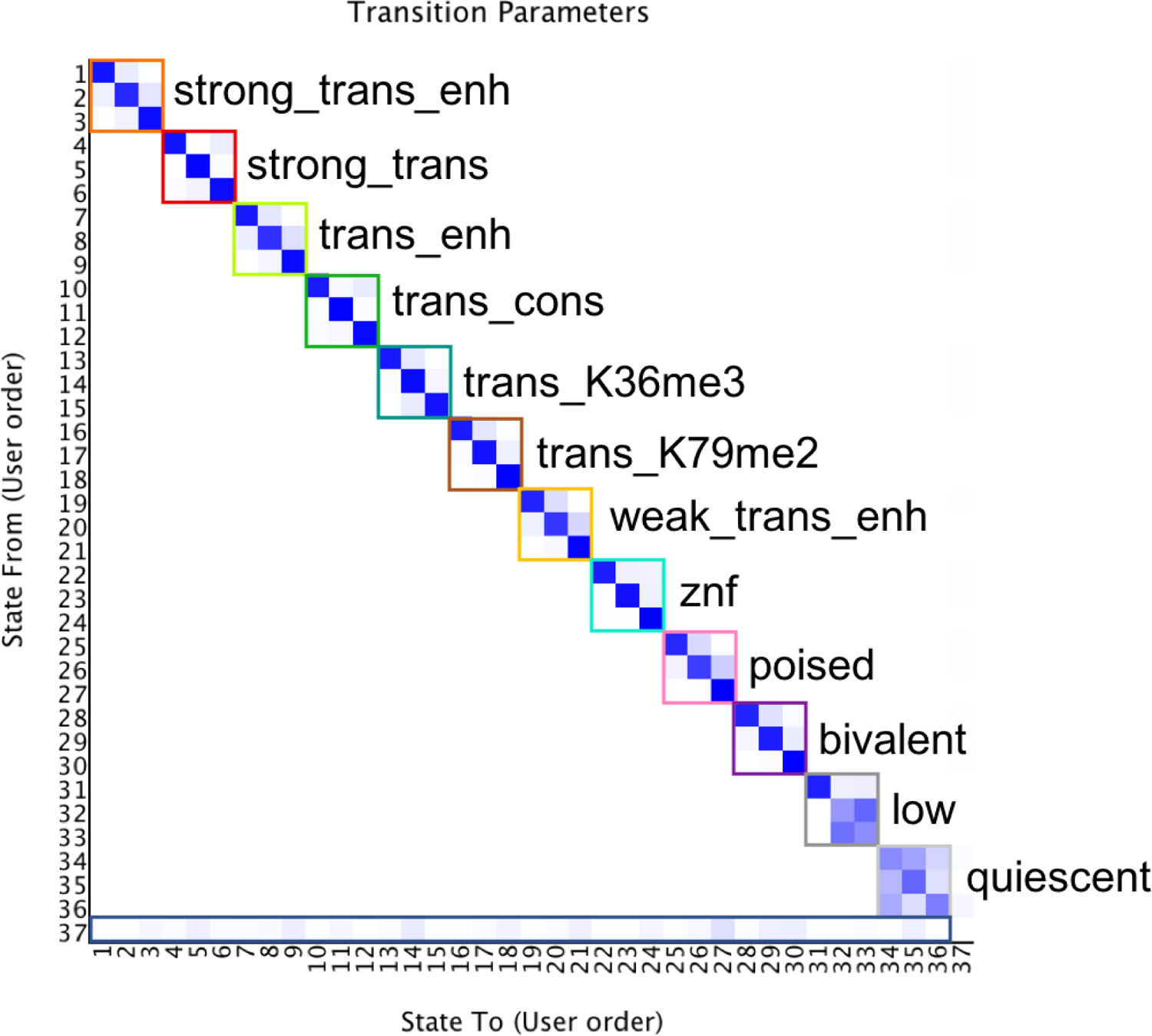
ChromGene state transitions Transition parameters between states within each component are marked in 3×3 squares. The bottom bar represents the initial probability of each state, independent of component. The component prior probabilities are the sums of the initial probabilities for their corresponding states. Transition probabilities are reported in (**Supp. Table 1**).

**Supplementary Figure 3:**
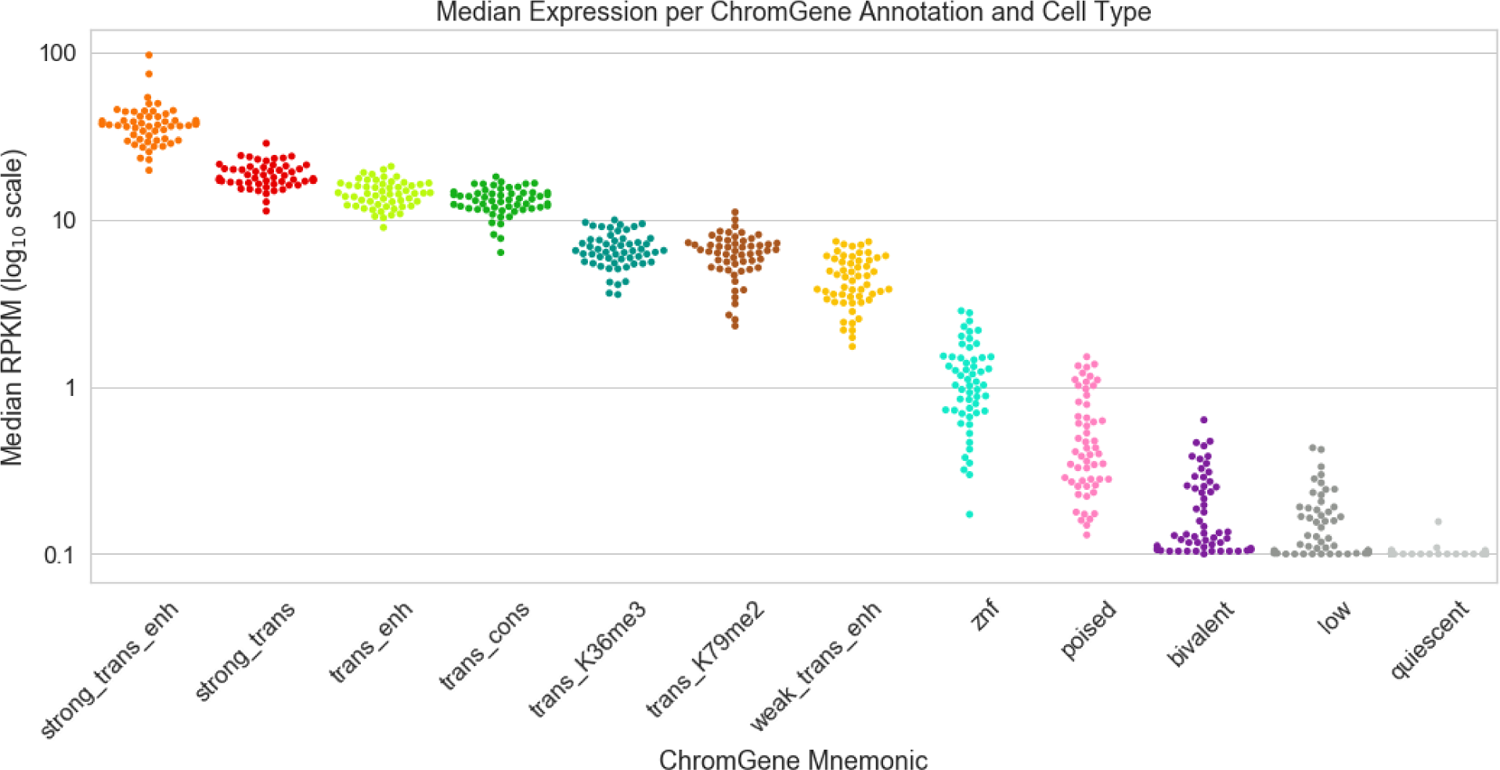
Median Expression for each ChromGene assignment, separated by cell type Distribution of median expression (RPKM) per ChromGene assignment over 56 cell types with expression available. Most points in the ‘quiescent’ annotation were 0.1, and not were plotted due to space. Median expression values were calculated based on all genes assigned to the ChromGene annotation in the cell type. Each point represents a single cell type and ChromGene assignment.

**Supplementary Figure 4:**
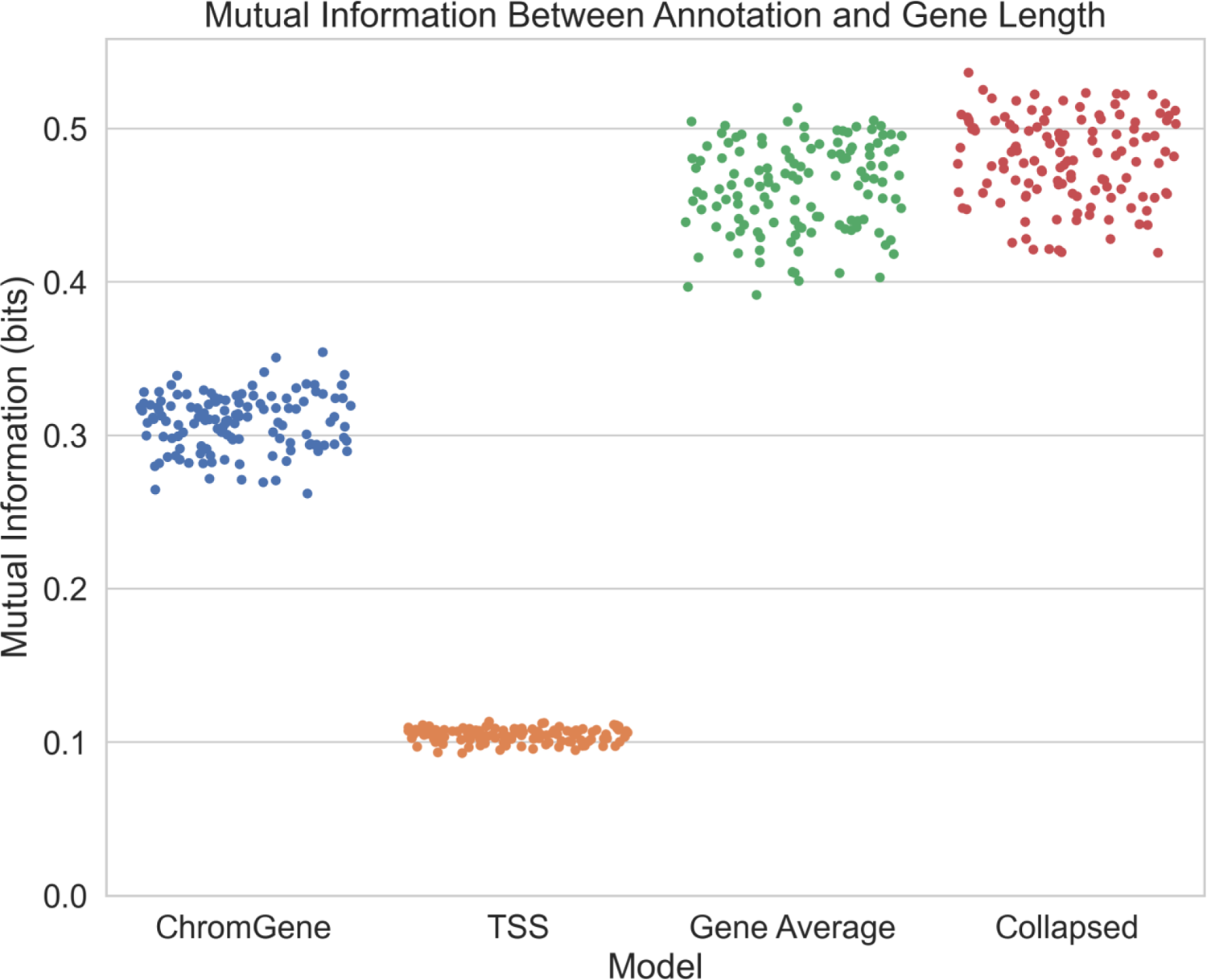
Mutual information of annotation and gene length The mutual information of annotations and gene lengths, where each point represents a cell type. Horizontal axis: ChromGene and three baselines: TSS, gene average, and collapsed models. ChromGene assignments have less information shared with the gene length (p < 10^-30^, paired binomial test, **Methods**), indicating that ChromGene annotations are less likely to directly reflect information about gene length compared to baseline methods that incorporate information from the whole gene. Annotations from the TSS model have the lowest mutual information with gene length, as expected, given it only uses information at the TSS.

**Supplementary Figure 5:**
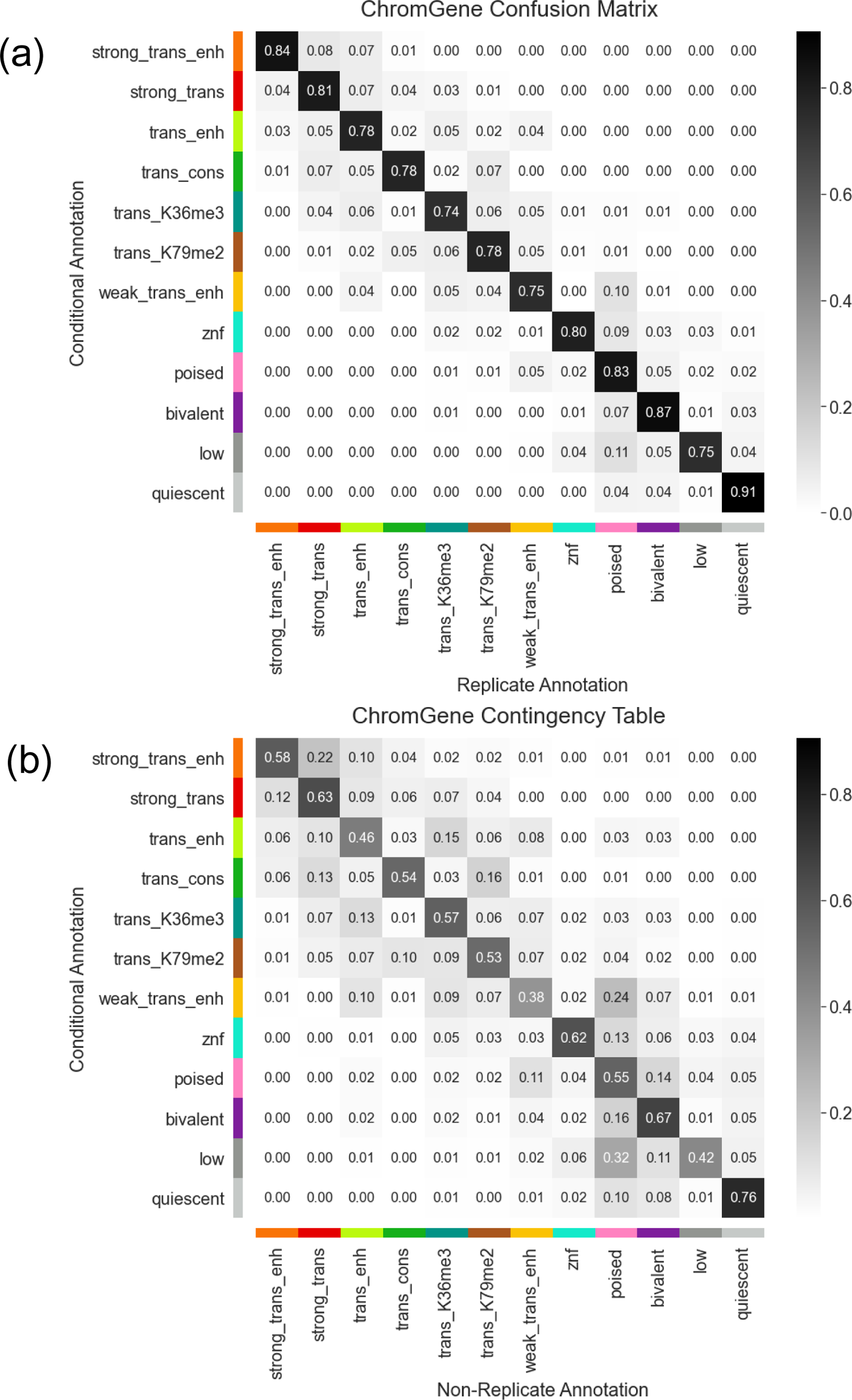
ChromGene confusion matrix and contingency table (a) Confusion matrix for ChromGene annotations between pairs of cell types that can be treated as biological replicates. (b) Contingency table of probabilities between pairs of cell types that are not biological replicates (**Methods**). In both the confusion and contingency matrices, rows correspond to the annotation being conditioned on and sum to unity probability.

**Supplementary Figure 6:**
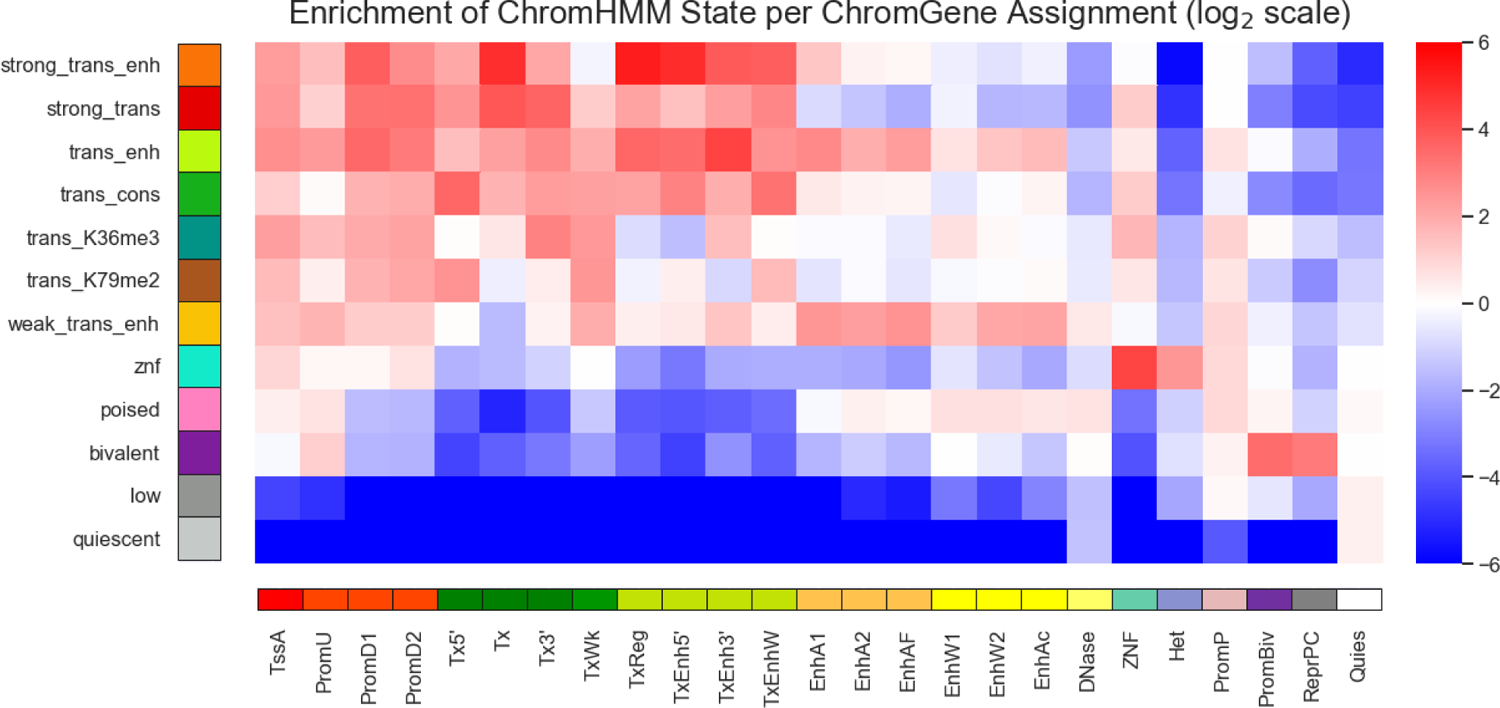
Log_2_ enrichments of ChromHMM states for each ChromGene assignment. Heatmap where the rows correspond to different ChromGene assignments and the columns to different cell type-specific chromatin states from a previous ChromHMM 25-state model based on the same imputed data [27]. Heatmap values indicate log_2_ enrichments for the matched chromatin state annotation in the ChromGene annotation. We set the minimum log_2_ enrichment to −6 to prevent washing out of the color scale due to highly negative values.

**Supplementary Figure 7:**
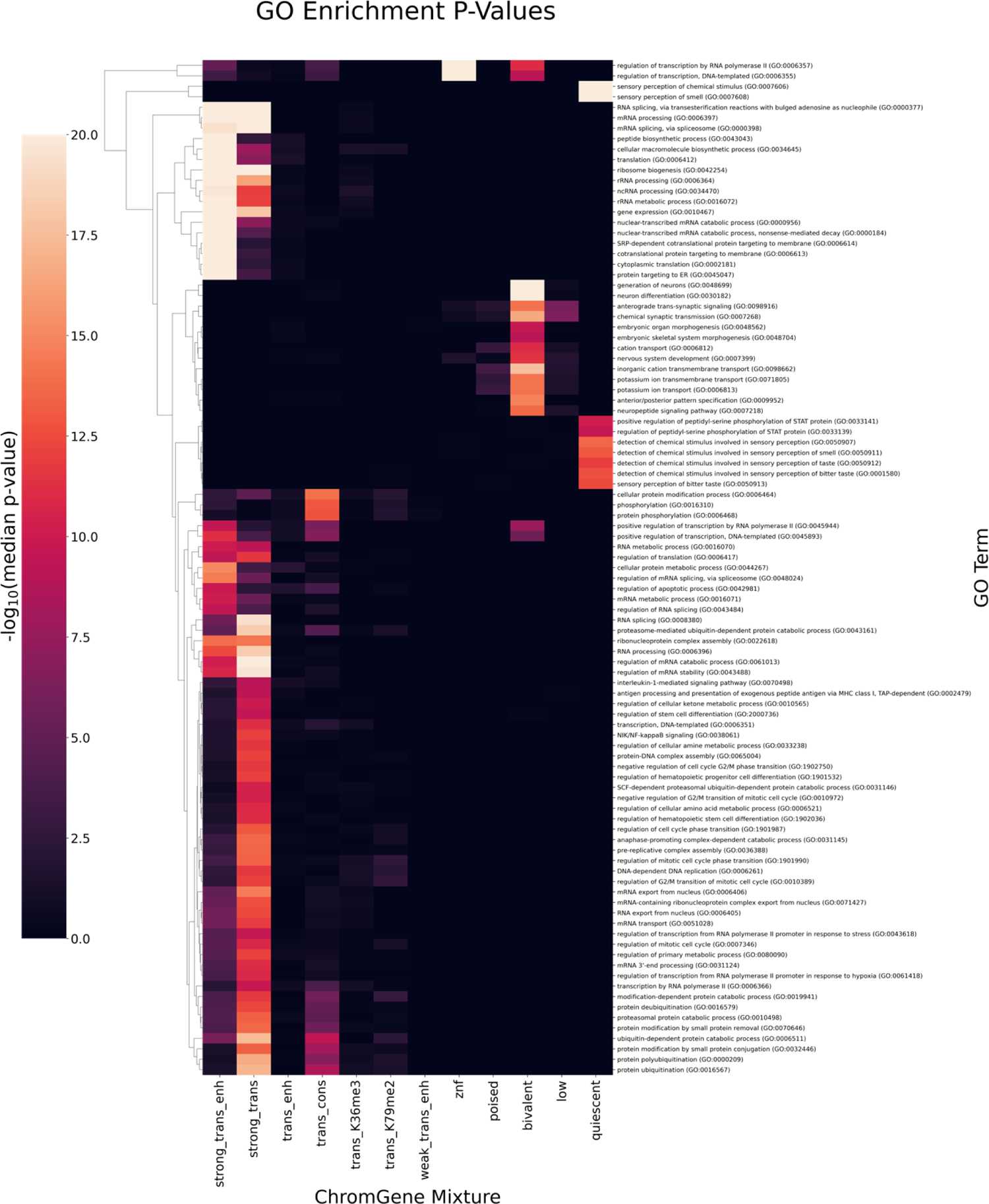
Median GO term enrichment across all cell types Heatmap where the row corresponds to GO terms, columns to ChromGene annotations, and values to the median enrichment -log_10_(p-value) across all cell types. Rows are filtered so that at least one ChromGene annotation is significant (adjusted p-value < 0.01), and are ordered using hierarchical clustering. Maximum -log_10_(p-value) is set to 20 to prevent washing out of the color scale.

**Supplementary Figure 8:**
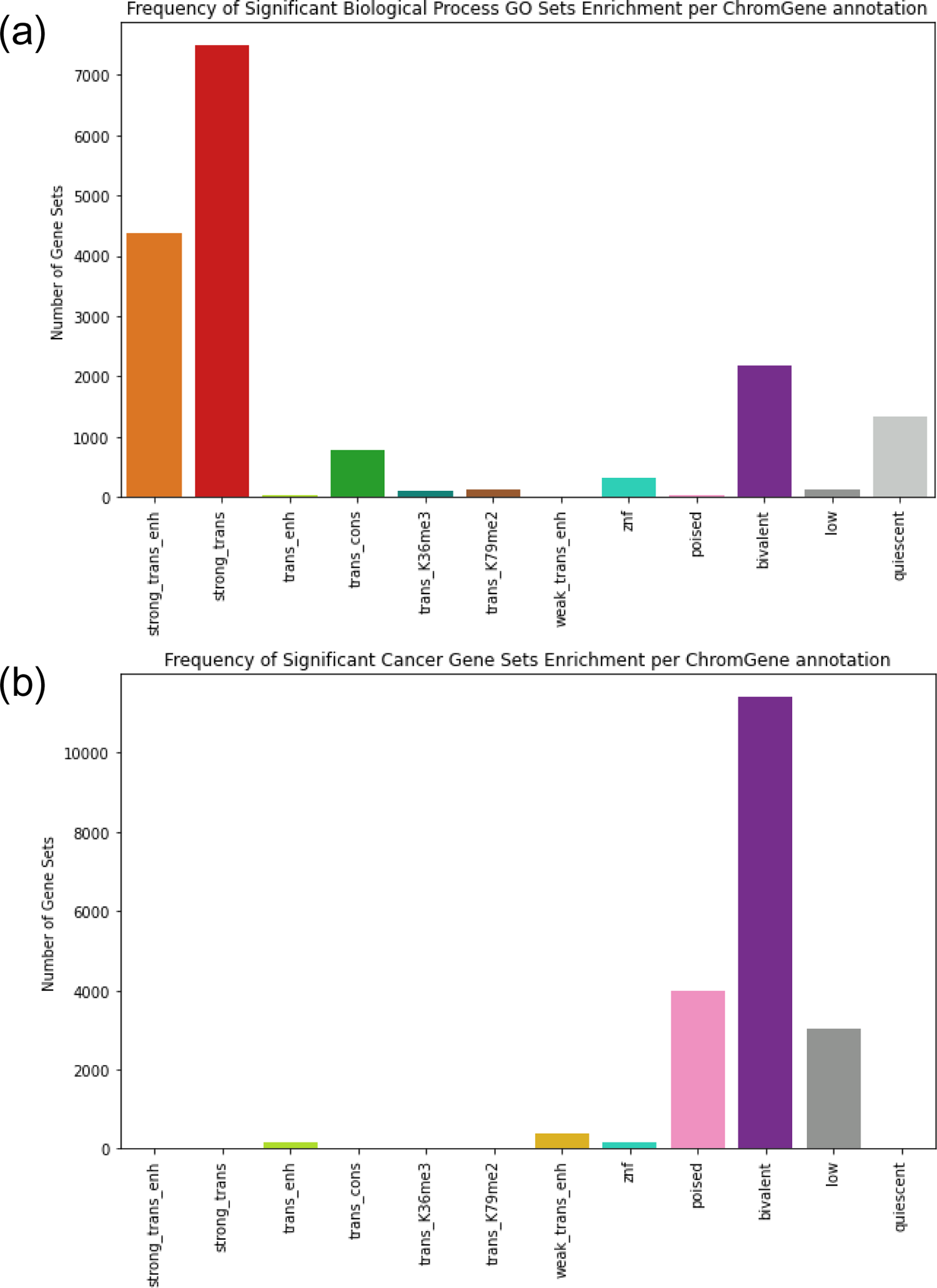
Frequency of ChromGene annotations for GO term and cancer gene set enrichments Count of how often each ChromGene annotations yielded significant p-values (p < 0.01, Bonferroni corrected for the number of combinations of cell types (127), components (12), and gene sets) across (a) 6036 gene sets for biological process GO terms and (b) 967 cancer gene sets.

**Supplementary Figure 9:**
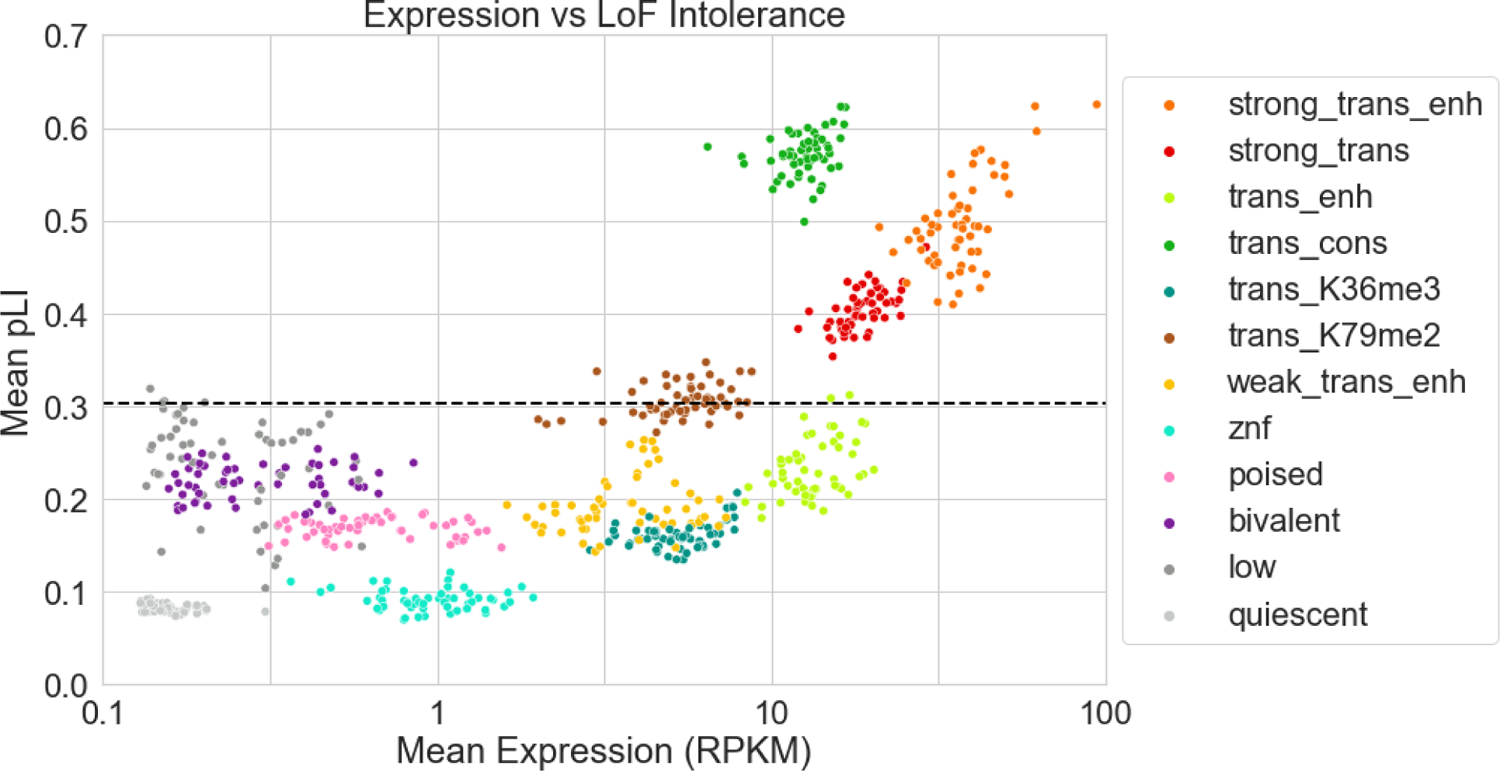
Mean pLI vs mean expression For each cell type and each ChromGene annotation combination, the mean RPKM+0.1 in log_10_ scale (x-axis) and mean pLI (y-axis) are plotted. Different ChromGene annotations are shown with different colors.

## Legends for additional supplementary files

**Supplementary Table 1:**

**Annotation enrichments tab –** The columns on this tab after the **Annotation** colors are as follows: Mnemonic: short identifying name of ChromGene annotation

Description: short description of ChromGene annotation

Overall percentage: percentage of gene-cell type combinations assigned to annotation

Median expression: median expression (RPKM) of genes assigned to annotation across 56 cell types with expression [1]

Median length (kb): median length (kb) of genes assigned to annotation across all cell types, not including flanking regions

Percentage of high-pLI genes (pLI > 0.9): percentage of genes across cell types that have a pLI score > 0.9 Contingency table diagonal / confusion matrix diagonal: percentage consistency of annotations across non-replicate cell types divided by percentage consistency of annotations across replicate cell types Cell type specificity: 1 - (contingency table diagonal / confusion matrix diagonal), a metric of cell type specificity

\# Housekeeping gene: gene annotated as housekeeping [35]

Housekeeping gene percentage: percentage of gene-cell type combinations annotated as a housekeeping gene

Housekeeping gene enrichment: fold enrichment of housekeeping genes compared to overall percentage Housekeeping gene log_2_ enrichment: log_2_ fold enrichment of housekeeping genes

Housekeeping gene enrichment median enrichment p-value: median p-value of housekeeping gene enrichment across cell types

\# Constitutively unexpressed gene: gene that has RPKM < 1 across 56 cell types with expression [1] Constitutively unexpressed gene percentage: percentage of gene-cell type combinations annotated as constitutively unexpressed

Constitutively unexpressed gene enrichment: fold enrichment of constitutively unexpressed genes compared to overall percentage

Constitutively unexpressed gene log_2_ enrichment: log_2_ fold enrichment of constitutively unexpressed genes Constitutively unexpressed gene median enrichment p-value: median p-value of constitutively unexpressed gene enrichment across cell types

\# Constitutively expressed gene: gene that has RPKM > 1 across 56 cell types with expression [1]

Constitutively expressed gene percentage: percentage of gene-cell type combinations annotated as constitutively expressed

Constitutively expressed gene enrichment: fold enrichment of constitutively expressed genes compared to overall percentage

Constitutively expressed gene log_2_ enrichment: log_2_ fold enrichment of constitutively expressed genes Constitutively expressed gene median enrichment p-value: median p-value of constitutively expressed gene enrichment across cell types

\# Olfactory gene: gene annotated as olfactory [30]

Olfactory gene percentage: percentage of gene / cell type combinations annotated as olfactory

Olfactory gene enrichment: fold enrichment of olfactory genes compared to overall percentage

Olfactory gene log_2_ enrichment: log_2_ fold enrichment of olfactory genes

Olfactory gene median enrichment p-value: median p-value of olfactory gene enrichment across cell types

\# ZNF gene: gene starts with “ZNF“

ZNF gene percentage: percentage of gene / cell type combinations annotated as ZNF

ZNF gene enrichment: fold enrichment of ZNF genes compared to overall percentage

ZNF gene log_2_ enrichment: log_2_ fold enrichment of ZNF genes

ZNF gene median enrichment p-value: median p-value of ZNF gene enrichment across cell types

Cancer gene sets enriched (adj p < 0.01): the number of cancer gene sets enriched across all cell types for given annotation

Cancer gene sets enriched percentage: the percentage of cancer gene sets enriched across all cell types for given annotation

BP GO terms enriched (adj p < 0.01): the number of ‘Biological Process’ GO term gene sets enriched across all cell types for given annotation

BP GO terms enriched percentage: the percentage of ‘Biological Process’ GO term gene sets enriched across all cell types for given annotation

Color (hex): hex color for ChromGene annotation

Matplotlib color name: color used in matplotlib for ChromGene annotation [https://xkcd.com/color/rgb/]

**State emissions and enrichments tab –** The first column gives a color and number for each annotation. The second column gives the annotation mnemonic. The third column gives a number to each individual state of the mixture component. The next 12 columns give the emission probabilities for each chromatin mark as indicated. The next two columns gives the maximum and minimum emission probabilities represented as percentages. The next column gives the enrichment of the individual states for annotated TSS. Individual states within each mixture component are ordered in decreasing value of this enrichment. The next column gives the initial probability of starting in the state overall. The last column gives the initial probability of the state given the component.

**Transition Probabilities tab –** This tab shows the transition probability indicating the probability when in the state of the row of transitioning to the state of the column. Probabilities are shown for individual states of the model which are ordered and colored based on the annotation to which they belong as indicated.

### Supplementary Data: ChromGene annotations

This is a gzip of a tab delimited file containing the ChromGene annotations for the 127 cell types. Each row after the header row corresponds to one gene. The first five columns from left to right are the chromosome of the gene, the left-most coordinate of the gene, the right-most coordinate of the gene, the gene symbol, and strand of the gene. This is based on hg19 and ENSEMBL v65/GENCODE v10. The remaining columns correspond to different cell types for which ChromGene annotations are reported as indicated by the names in the header.

